# Young children rely on visual information to process degraded speech: Evidence from behavioural and neuroimaging measures

**DOI:** 10.64898/2026.03.05.707650

**Authors:** Irene Arrieta-Sagredo, Borja Blanco, César Caballero-Gaudes, Manuel Carreiras, Marina Kalashnikova

## Abstract

Young children acquire language in environments where speech is often acoustically degraded, yet little is known about how developing brains adapt to reduced speech intelligibility. Using a combination of eye-tracking and functional near-infrared spectroscopy (fNIRS), we investigated young children’s attentional allocation to a speaking face at varying levels of speech intelligibility and the brain activity supporting this behaviour during development. Infants (8–10 months) and toddlers (27–30 months) viewed videos of a speaker in three conditions: producing clear speech, spectrally degraded (vocoded) speech, and silent (audio muted) speech. Visual attention to the speaker’s mouth increased when speech was degraded relative to clear speech in both age groups, indicating an early-emerging compensatory strategy. However, this shared behavioural response was supported by brain activity that differed by age. Degraded speech elicited greater recruitment of prefrontal regions associated with effortful listening, particularly in infants, whereas toddlers showed stronger engagement of posterior temporal regions implicated in audiovisual integration. In response to silent speech, there was no evidence for increased visual attention to the mouth compared to the clear speech condition, but there was reduced temporal activation and increased prefrontal brain responses, especially in infants. Together, these findings suggest that experience with audiovisual correspondences and linguistic maturity contribute to a more efficient processing of speech, particularly relevant when speech is degraded. By combining behavioural and neuroimaging measures, this study advances mechanistic accounts of audiovisual speech processing and provides insights relevant to populations experiencing spectrally degraded input, such as children using cochlear implants.

## 1 Introduction

Spoken language often occurs in challenging acoustic conditions, when understanding is reduced by environmental factors such as background noise and competing talkers (Sumby & Pollack, 1954; Vatikiotis-Bateson et al., 1998). Under these circumstances, attention to visual cues from a speaker’s face substantially enhances comprehension in adults (Chandrasekaran et al., 2009; Van Engen et al., 2022). This audiovisual benefit reflects the integration of redundant and complementary information across modalities, allowing listeners to compensate for deficient auditory input. Infants and young children also acquire language in imperfect listening environments, but less is known about the ability to utilise audiovisual information during the first years of life, and how maturation and growing language experience shape the brain activity that supports it.

Visual attention to the speaker’s face is a behavioural marker of audiovisual speech processing. During the first months of life, infants direct greater attention to speakers’ eyes than to the mouth in response to familiar continuous infant-directed speech. This is attributed to the eyes’ social salience (Santapuram et al., 2022) and physical contrast (Otsuka et al., 2013). Around 8 to 10 months of age, a shift of attention towards the mouth takes place (Chawarska et al., 2009; Hillairet de Boisferon et al., 2018; Lewkowicz & Hansen-Tift, 2012; Mercure et al., 2019; Morin-Lessard et al., 2019; Pons et al., 2015; Sekiyama et al., 2021; Tenenbaum et al., 2014), and it is maintained until approximately the second year of age, when a more balanced distribution of attention between the eyes and the mouth emerges (Morin-Lessard et al., 2019; Shic et al., 2020). Attention to the mouth allows infants access to visual cues that support speech processing, especially at the time when their own speech perception and production abilities undergo significant development, coinciding with important milestones, such as the onset of canonical babbling, vocal imitation (Imafuku et al., 2019; Imafuku & Myowa, 2016), and phonological learning (Lewkowicz & Hansen-Tift, 2012). Indeed, individual differences in the amount of attention directed to a speaker’s mouth have been positively correlated with expressive vocabulary size (Chawarska et al., 2009; Frank et al., 2012; Hillairet de Boisferon et al., 2018; Nakano et al., 2010; Shic et al., 2020; Tenenbaum et al., 2014), word recognition (Król, 2018), and grammatical rule learning (Birulés et al., 2022). On the other hand, children with developmental language disorders show reduced attention to the mouth (Pons et al., 2018), underscoring its developmental relevance.

Importantly, experienced language users can deploy visual attention to the mouth when speech is difficult to understand (Banks et al., 2021; Vatikiotis-Bateson et al., 1998), but when this compensatory strategy is developed and its role in facilitating language processing early in life are unknown. There is evidence that 10-month-old infants look longer to the mouth of a speaker of an unfamiliar than a familiar language (Birulés et al., 2023; Lewkowicz & Hansen-Tift, 2012; Morin-Lessard et al., 2019; Pons et al., 2015). Bilingual infants, who face the additional challenge of differentiating their two languages during speech processing, start to look longer at a speaker’s mouths already by 4 months of age, earlier than their monolingual counterparts (Pons et al., 2015). To our knowledge, only one study directly investigated young children’s visual attention in response to speech with decreased intelligibility. Król (2018) presented 17-to-35-month-old toddlers with faces producing nursery rhymes, and intelligibility was reduced by creating an audiovisual mismatch for a proportion of syllables in the stimuli. Toddlers also completed an experimental measure of lexical knowledge (a word recognition task). As expected, toddlers directed greater attention to the speaker’s mouth in response to the less intelligible stimuli, but this was positively correlated with their word recognition performance. Thus, toddlers with more advanced lexical competence showed a greater reliance on visual information in noisy stimuli, as a strategy to access linguistic content and greater motivation to improve comprehension (Król, 2018).

Together, these findings suggest that young children can selectively attend to visual speech, and in the case of challenging listening conditions, the ability to use this strategy may emerge with growing language competence. However, it remains unknown when this compensatory mechanism emerges in infancy, especially in response to acoustically degraded speech. In addition, this evidence remains largely behavioural, leaving open the question of which neural mechanisms support this compensatory strategy during development. In adults, reduced speech intelligibility engages a distributed network of brain regions associated with effortful listening and audiovisual integration. Degraded speech elicits increased activity in frontal and cingulo-opercular regions involved in cognitive control, alongside changes in temporal regions supporting speech processing (Davis & Johnsrude, 2007; Peelle, 2018b; Wild et al., 2012). Audiovisual speech additionally recruits posterior superior temporal regions implicated in multisensory integration, as well as occipital and premotor areas that provide complementary visual and articulatory information (Beauchamp, 2016a; Erickson et al., 2014). Crucially, studies combining eye-tracking and neuroimaging show that attention to the mouth modulates activity in these networks, particularly under degraded listening conditions (Jiang et al., 2016).

In infants, unmodified, clear speech elicits responses in early cortical regions such as the primary auditory cortex, which are sensitive to the acoustic form of speech (Blanco et al., 2025; Dehaene-Lambertz et al., 2006, 2010; Dehaene-Lambertz & Spelke, 2015), whereas frontal, peri-auditory and premotor regions contribute to a more abstract processing of the linguistic stimuli (Altvater-Mackensen & Grossmann, 2016, 2018; Dopierała et al., 2023; Mercure et al., 2020). In response to audiovisual speech, similar mechanisms to those of adults may emerge early in development. For instance, Altvater-Mackensen and Grossmann (2016) presented 6-month-old infants with speaking faces producing vowels in conditions where the visual and auditory information was matched vs. mismatched. Brain responses, measured via functional near-infrared spectroscopy (fNIRS), revealed a trend for increased activation in the prefrontal, anterior cortex in the mismatched conditions. Moreover, there was a relation between brain activity in the prefrontal brain areas and infants’ looking time to the mouth of a speaking face (measured in a separate task). These findings were interpreted as reflecting the involvement of the inferior frontal gyrus in audiovisual integration, a region thought to play a role in linking and representing phonological speech information across modalities. However, this interpretation remains tentative, as brain activity in auditory and audiovisual cortical regions was not measured (Altvater-Mackensen & Grossmann, 2016). Moreover, it remains largely unknown whether similar neural correlates support attention to visual speech cues during naturalistic, continuous speech, and how these correlates change with maturation and growing language experience.

### The present study

As discussed above, young infants use the visual information from a speaker’s mouth to support speech processing. With increasing language experience, they develop the ability to selectively deploy attention to the mouth when listening conditions are challenging, and the use of this strategy may be subserved by the maturation of neural processes related to effortful listening and audiovisual integration, as it is the case in adults. The present study aimed to define the developmental trajectory for this attentional strategy and related brain activity.

For this purpose, we measured attentional allocation to a speaking face during continuous speech at varying levels of intelligibility, while concurrently measuring the brain activity supporting this behaviour in 9-month-old infants and 2.5-year-old toddlers. We chose these specific age groups as they correspond to the initial significant increase in attention to speakers’ mouth over the eyes and its subsequent decrease in favour of balanced attention to the mouth and the eyes in response to clear continuous speech (Lewkowicz & Hansen-Tift, 2012; Morin-Lessard et al., 2019; Pons et al., 2015). Furthermore, these age groups allowed us to test how the selective use of this strategy to support the processing of degraded speech is shaped by children’s access to linguistic information in the stimuli, whereby infants have more limited access to linguistic information that can support processing and comprehension compared to toddlers.

We presented children with videos of continuous infant-directed speech presented in three auditory conditions with progressively reduced auditory speech information: clear (unmodified), vocoded (8-channel vocoded, spectrally degraded), and silent (no auditory signal). Visual attention to the mouth versus eyes was measured using eye-tracking. Simultaneously, brain activity was recorded via functional near-infrared spectroscopy (fNIRS), targeting prefrontal, temporal, motor, and occipital regions. As explained below, the clear and vocoded conditions comprised the key comparison for this study, as these conditions both contained auditory and visual information but differed in the amount of speech intelligibility. The visual-only speech condition was also included with the secondary purpose of identifying the brain signatures of visual speech processing, to better understand the additive effects of auditory and visual information when measuring brain responses associated with the clear and vocoded audiovisual conditions. Additionally, this condition could provide relevant insights into visual speech processing in infants with very limited or no access to auditory information, such as infants born with hearing loss.

With regards to the key comparison, for the behavioural measures of visual attention, we hypothesized that decreasing speech intelligibility would be associated with increased visual attention to the mouth, reflecting greater reliance on visual speech cues and increased processing effort under more challenging listening conditions (Vatikiotis-Bateson et al., 1998). This effect could be observed across both infants and toddlers. Alternatively, we hypothesized that the relationship between intelligibility and visual attention would be moderated by age, such that increased attention to the mouth under low intelligibility conditions would be evident primarily in toddlers. This age-specific modulation would be consistent with toddlers’ more advanced linguistic processing abilities and greater motivation to allocate attentional resources to speech comprehension compared to infants (Król, 2018).

Second, we expected that the neural measures would reflect the hypothesised behavioural patterns. Specifically, we predicted that reduced speech intelligibility would be associated with increased brain activity in regions implicated in attention and effortful listening, particularly the prefrontal cortex (Davis & Johnsrude, 2003; Mushtaq et al., 2021; Peelle, 2018a), alongside decreased activation in temporal regions surrounding the primary auditory cortex, as reported in older children under degraded listening conditions (Mushtaq et al., 2021). These brain activation effects were expected to occur across age groups. Alternatively, if intelligibility effects are developmentally modulated, toddlers were expected to show greater increases in brain activation within regions supporting audiovisual integration, such as the posterior temporal cortex, under low intelligibility conditions. Finally, we conducted an exploratory analysis of occipital and motor regions, as these areas have been implicated in audiovisual speech processing in adults (Peelle, 2018a), but remain largely unexplored in infants in response to degraded audiovisual speech.

## 2 Method

### 2.1. Participants

A total of 64 children were recruited for the study. Twenty-one participants were excluded due to experimenter error, excessive fussiness or crying, or poor data quality (11 infants and 10 toddlers). The final sample comprised 43 children: 23 infants (mean age = 8.97 months, SD = 1.40; 6 females) and 20 toddlers (mean age = 29.00 months, SD = 4.38; 11 females). All 43 participants provided usable gaze data. In addition, usable fNIRS data were obtained from 18 infants (4 females) and 18 toddlers (9 females; see Section 2.5 for details). All children were typically-developing with no known hearing or vision deficits or risk for developmental delays.

All children were born and lived in the Basque Country in Northern Spain, a bilingual region where Basque and Spanish are official languages. Basque is the official language in early childhood education settings, but children also receive exposure to Spanish in the community. For these reasons, the experiment was conducted in Basque, and children with predominant exposure to Basque were included (i.e., more than 60% Basque exposure). To determine this, parents completed a Language Exposure Questionnaire in which they reported their children’s patterns of exposure to Spanish and Basque from birth, their own language proficiency levels in Spanish and Basque, and patterns of language use in the home (Bosch & Sebastián-Gallés, 2001). Infants’ Basque exposure ranged from 77.4% to 100% (mean = 93.8%, SD = 7.8%), and toddlers’ from 62.4% to 100% (mean = 87%, SD = 13.9%).

Caregivers signed an informed consent form before starting the experimental procedure. The study was approved by the Basque Center on Cognition, Brain and Language (BCBL) Ethics Committee and conducted in accordance with the relevant guidelines and regulations. No monetary compensation was offered to the families, but a small gift was given to the children at the end of the experimental session.

### 2.2. Apparatus: Eye-tracker and fNIRS

Eye gaze was acquired using an EyeLink 1000 eye-tracker and Experimental Builder software. Stimuli were presented on screen (34-27 cm), separated around 70 cm from the infant (Figure 1A). The audio was presented at 65-70 dBA SPL through two free-field JBL loudspeakers. Simultaneously, brain activity was measured with a continuous wave fNIRS system (NIRScout), consisting of three components: 16 optodes (laser emitting diodes on two different wavelengths: 760nm and 850 nm), 16 detectors and a control box. Optodes were mounted on a custom-built headgear (actiCAP) comprising 40 measurement channels with a 3-cm source–detector separation. The montage covered prefrontal, premotor, temporal, and occipital cortical regions and was positioned according to the international 10–20 system (Figure 2B). Data were acquired at a sampling rate of 3.9 Hz using NirStar 15.3 software.

**Figure 1.**
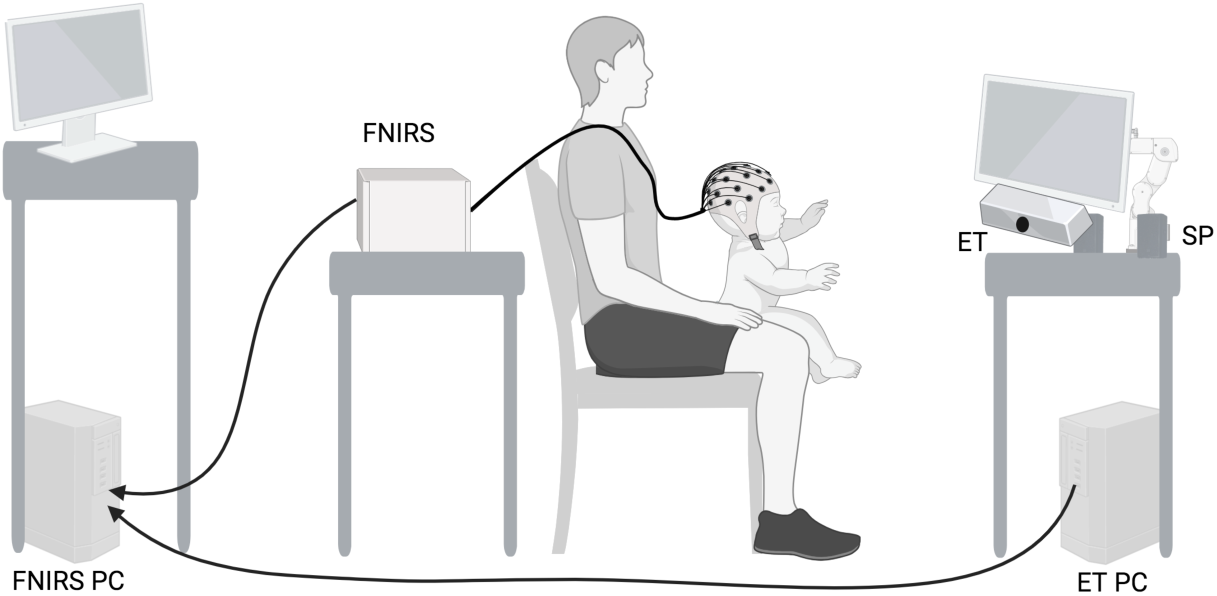
Schematic representation of the experimental setup (created using BioRender).

**Figure 2.**
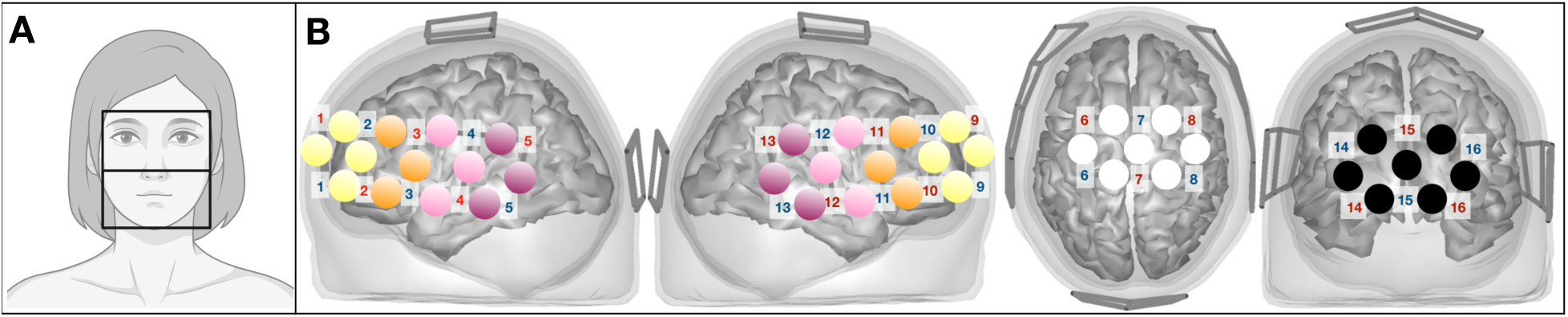
Regions of interest (ROIs) used in the analyses. (A) Eye-tracking ROIs for gaze analysis (eyes and mouth, note that this is just an schematic illustration, we used a real female video). (B) fNIRS ROIs: temporal cortex ROI, subdivided into posterior temporal (purple) and anterior temporal (pink) regions; prefrontal cortex ROI, subdivided into posterior prefrontal (orange) and anterior prefrontal (yellow) regions; motor cortex ROI (white); and occipital cortex ROI (black). Sources (red) and detectors (blue) are displayed with identification numbers overlaid on the montage, and channels are color-coded according to ROI.

### 2.3. Stimuli

A female native speaker of Basque was recorded producing three brief passages in infant-directed speech. The passages were comparable in duration (16–18 s each) and each consisted of eight short utterances (see Supplementary Material 1). The speaker was recorded on video showing her full face and part of the neck against a white background (Figures 1B and 2A). At the beginning and end of each recording, the speaker appeared smiling for one second.

The original video recordings with clear, unprocessed audio were used for the Clear condition. The Vocoded condition was created by dividing the clear audio into eight spectral bands and applying their amplitude envelopes to white noise (with equalised loudness curves), preserving coarse amplitude modulation while replacing fine spectral structure with noise (Shannon et al., 1995). Eight spectral bands were chosen because this number is commonly used to simulate cochlear implant perception (Cabrera & Werner, 2017; Hegde et al., 2024). The Silent condition consisted in removing the audio signal from the video. As a result, nine videos were created for the experimental trials, three for each condition.

### 2.4. Procedure

Children sat on their caregiver’s lap, wearing the fNIRS cap and facing the screen mounted on top of the eye-tracking camera (see Figure 1A). First, fNIRS calibration was completed (allowing for a maximum of 4 attempts to obtain good or medium calibration on as many channels as possible), followed by a five-point calibration on the eye-tracker. The eye-tracking and fNIRS signals were co-registered with the eye-tracking software (Experimental builder) by sending a trigger to the fNIRS system (NIRScout) every time a trial started and ended.

After calibration, children were first presented with 9 experimental trials, 3 passages in 3 conditions each (clear, silent, and vocoded). An attention getter (a silent display of colourful stars on a black screen) with duration jittered between 3 and 6 seconds was presented before the start of each trial. Children first saw a silent trial, a clear trial, and a vocoded trial. This fixed initial sequence was chosen to promote engagement during the silent condition and to reduce surprise effects associated with the acoustically unnatural vocoded speech. The remaining 6 trials were presented in one of six experimental orders (consisting of different pairings between passages and conditions and a different order of trials belonging to the three conditions, see Supplementary Material 2) counterbalanced across participants. After the 9 trials were completed, a single point drift correction was applied to make sure that the eye gaze calibration was correct. If the drift correction was correct, the child was presented with a repetition of the 9 trials. If the drift correction was not correct, eye gaze calibration was performed again before continuing. Thus, the task consisted of 18 trials in total and lasted for 6 minutes on average.

### 2.5. Data processing and analysis

#### 2.5.1. Pre-processing of gaze data

Gaze data were pre-processed using the *eyetrackingR* toolbox version 0.2.1 (Forbes et al., 2023) and custom scripts in R version 4.2.3 (R Core Team, 2021). Data for individual trials were excluded if participants fixated the screen for less than 60% of the trial. A participant’s data were included in the analysis if they contributed at least two valid trials in at least one experimental condition (Clear, Vocoded, Silent). After applying these criteria, the final sample consisted of 23 infants (out of 34 tested) and 20 toddlers (out of 30 tested). Detailed information on data loss per participant and condition is provided in Supplementary Materials 3 and 4.

Two regions of interest (ROIs) of equal dimensions covering the top half (eyes) and bottom half (mouth) of the speaker’s face were selected (see Figure 2A). For each trial, the proportion of total looking time to the eyes (PTLT eyes) was subtracted from the total looking time to the mouth (PTLT mouth). Next, the PTLT difference was normalised by dividing it by the total looking time to the face (Lewkowicz & Hansen-Tift, 2012). Scores > 0 represent greater fixation to the mouth than eyes, and scores < 0 greater fixation to the eyes than mouth.

#### 2.5.2. Statistical analysis of gaze data

Data were analysed using a linear mixed-effects model fitted using the *lme4* package in R (Bates et al., 2015) to examine attention to the speaker’s mouth relative to the eyes (dependent variable) across Group (infants, toddlers), Condition (Clear, Vocoded, Silent), and their interaction (independent variables). Subject was entered as a random intercept.

#### 2.5.3. Pre-processing of brain data

fNIRS data were processed using the *nirs-toolbox* in Matlab (Santosa et al., 2018). Channels were grouped into three ROIs across the two hemispheres based on the AAL atlas, choosing a specificity of 30% (default) (Fu & Richards, 2021): fronto-temporal, motor, and occipital (see Figure 2A and Supplementary Material 3 for more details about channel assignment to each ROI). Given the spatial resolution of fNIRS, anatomical labels refer to approximate cortical regions rather than the precise functional localization.

Only participants who met the eye-tracking inclusion criteria were considered for fNIRS data analyses, ensuring that only infants who attended to the task were included. fNIRS data quality was assessed at both the channel and trial levels using established signal quality metrics implemented in the *nirs-toolbox*.

At the channel level, channels were included if they demonstrated adequate optode-scalp coupling above 0.7 (Pollonini et al., 2016). At the trial level, trials were considered valid if the cardiac pulse was present for at least one third of the trial duration (Lee, Mao, Savkovic, et al., 2023) (see Supplementary Material 5). Participants were included if they retained more than 50% of good-quality channels overall, had at least two valid channels within the temporal ROI, and contributed a minimum of two valid trials per condition. After applying these criteria, the final sample for the fNIRS analyses consisted of 18 infants (out of 23 with valid gaze data) and 18 toddlers (out of 20 with valid gaze data). Full details of channel- and trial-level quality assessment procedures, thresholds and data loss are provided in Supplementary Materials 3, 4, 5 and 6.

Once quality assessment was performed, the raw intensity signals were converted to optical density. Next, motion artefact correction was performed by applying the temporal derivative distribution repair (TDDR) as part of the *nirs-toolbox* (Santosa et al., 2018). This algorithm decreases the amplitude of time points with big amplitude changes by decreasing the weight of those time points whose derivative exceeds a threshold to be considered biologically plausible (Fishburn et al., 2019). Signals were then band-pass filtered from 0.01 to 0.25 Hz (to avoid interference from physiological signals such as respiratory (∼0.3 Hz) and cardiac (∼>1 Hz) signals). Optical density signals were converted into changes in haemoglobin concentration using the modified Beer Lambert Law (Cope et al., 1988).

#### 2.5.4. Statistical analysis of brain data

fNIRS data were analysed including the estimation of the amplitude of the hemodynamic response at each channel for each participant as the dependent variable and Condition, Group and their interaction as independent variables (subject as a random effect).

The estimated hemodynamic response was generated by convolving a hypothesized stimulus time course based on the timings of the experimental trials with a canonical hemodynamic response function adapted to the 18-s stimulus duration. The comparison between the hypothesized and measured hemodynamic responses (see the measured hemodynamic time series in Supplementary Material 7) was performed using a general linear model (GLM) that yielded beta coefficients reflecting the correspondence between hypothesized and measured hemodynamic responses. Given that fNIRS records at a faster sampling frequency than the slow physiological signals recorded, the probability of measuring serially correlated signals is high, thus increasing the probability of Type I errors. To account for this, GLM parameter estimation employed an autoregressively whitened weighted least-squares regression model (AR-IRLS), which includes a pre-whitening step to remove temporal dependencies between successive samples.

At the group level, subject-level models were incorporated into group-level analyses for which the model was whitened using the error covariance of the first-level GLM (Santosa et al., 2018). Then, a correction for multiple comparisons was performed, using Benjamini-Hochberg FDR-corrected values (q<0.05) (Santosa et al., 2018).

## 3 Results

### 3.1. Visual attention to the speaker’s face

Only the main effect of Condition was statistically significant (see Figure 3 and Table 1 for full model results). Post-hoc pairwise comparisons indicated significantly lower PTLT to the mouth in the Clear condition compared to the Vocoded condition, t(395) = −2.718, p = .018. No significant differences were observed between the Clear and Silent conditions, t(396) = −0.319, p = .945, or between the Vocoded and Silent conditions, t(395) = 2.271, p = .061.

**Figure 3.**
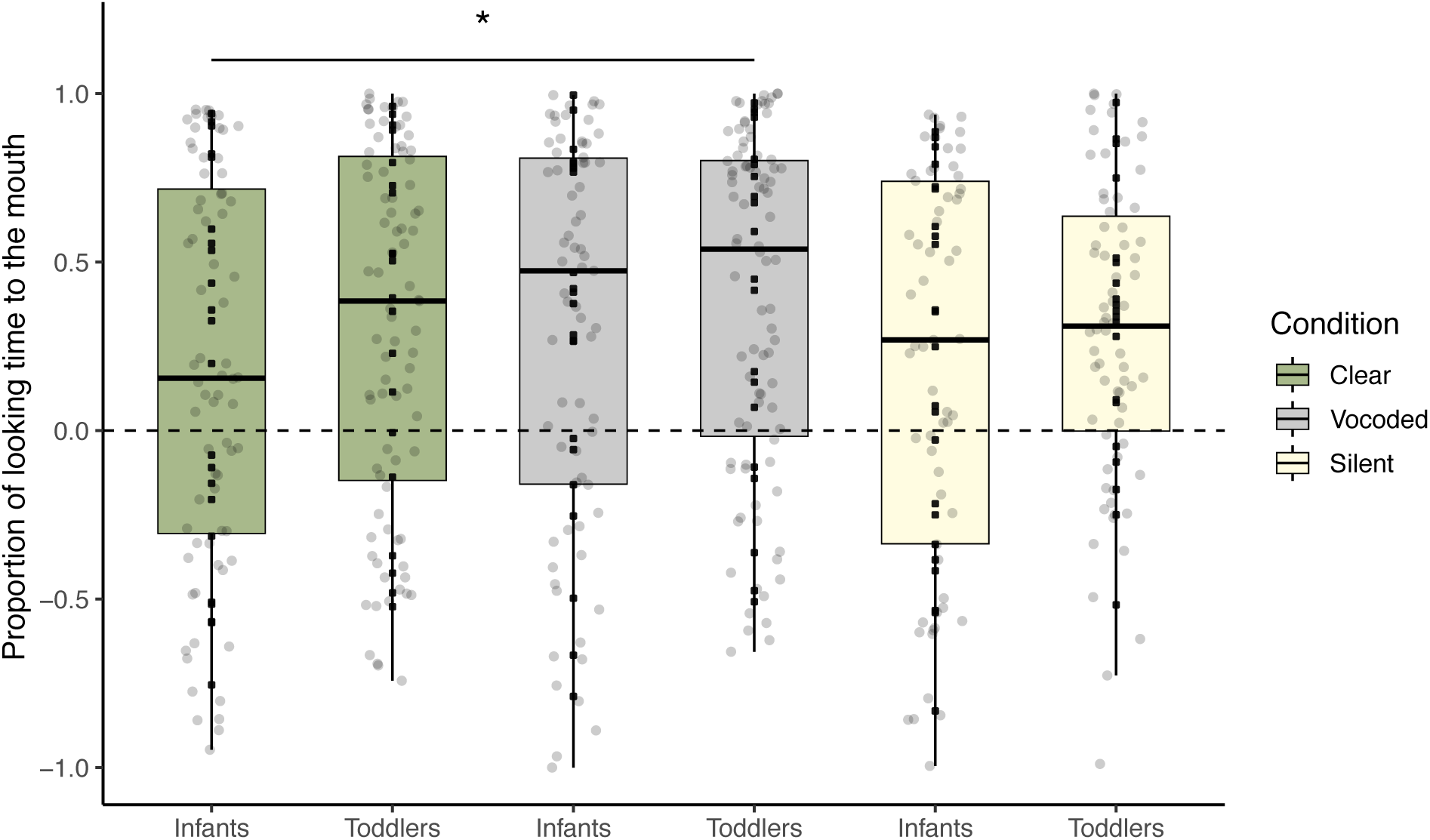
Proportion of total looking time (PTLT) directed to the mouth relative to the eyes in infants and toddlers across the clear, vocoded, and silent conditions. Grey dots represent individual trials, and black dots represent mean PTLT values per participant.

**Table 1.** Analysis of variance results for fixed effects from the linear mixed-effects model predicting the difference in proportion of total looking time to the mouth versus the eyes as a function of Condition (clear, vocoded, silent), Group (infants, toddlers), and their interaction, with a random intercept for Subject.

|  | Sum Sq | Mean Sq | NumDF | DenDF | F value | Pr (>F) |
| --- | --- | --- | --- | --- | --- | --- |
| Group | 0.008 | 0.008 | 1 | 41.06 | 0.535 | .468 |
| Condition | 0.143 | 0.071 | 2 | 395.25 | 4.323 | .013* |
| Group:Condition | 0.008 | 0.004 | 2 | 395.25 | 0.254 | .775 |

### 3.2. Brain responses

#### 3.2.1. Main effect of Age group on neural responses

A main effect of group was observed in the channels plotted in Figure 4. One channel on the anterior part of the left prefrontal cortex and two channels located on the right temporal hemisphere revealed higher HbO in infants compared to toddlers. On the contrary, one channel on the motor cortex showed higher HbO in toddlers compared to infants. For HbR, toddlers showed a more positive beta value than infants in two channels located on the left anterior prefrontal cortex, on one channel at the right posterior temporal cortex, and on one channel at the right anterior prefrontal cortex. However, from the reported results, only one channel for HbR, located in the right anterior prefrontal cortex (marked inside a black box in Figure 4), survived the correction for multiple comparisons using the Benjamini-Hochberg FDR-corrected values (*q* < .05) (Santosa et al., 2018).

**Figure 4.**
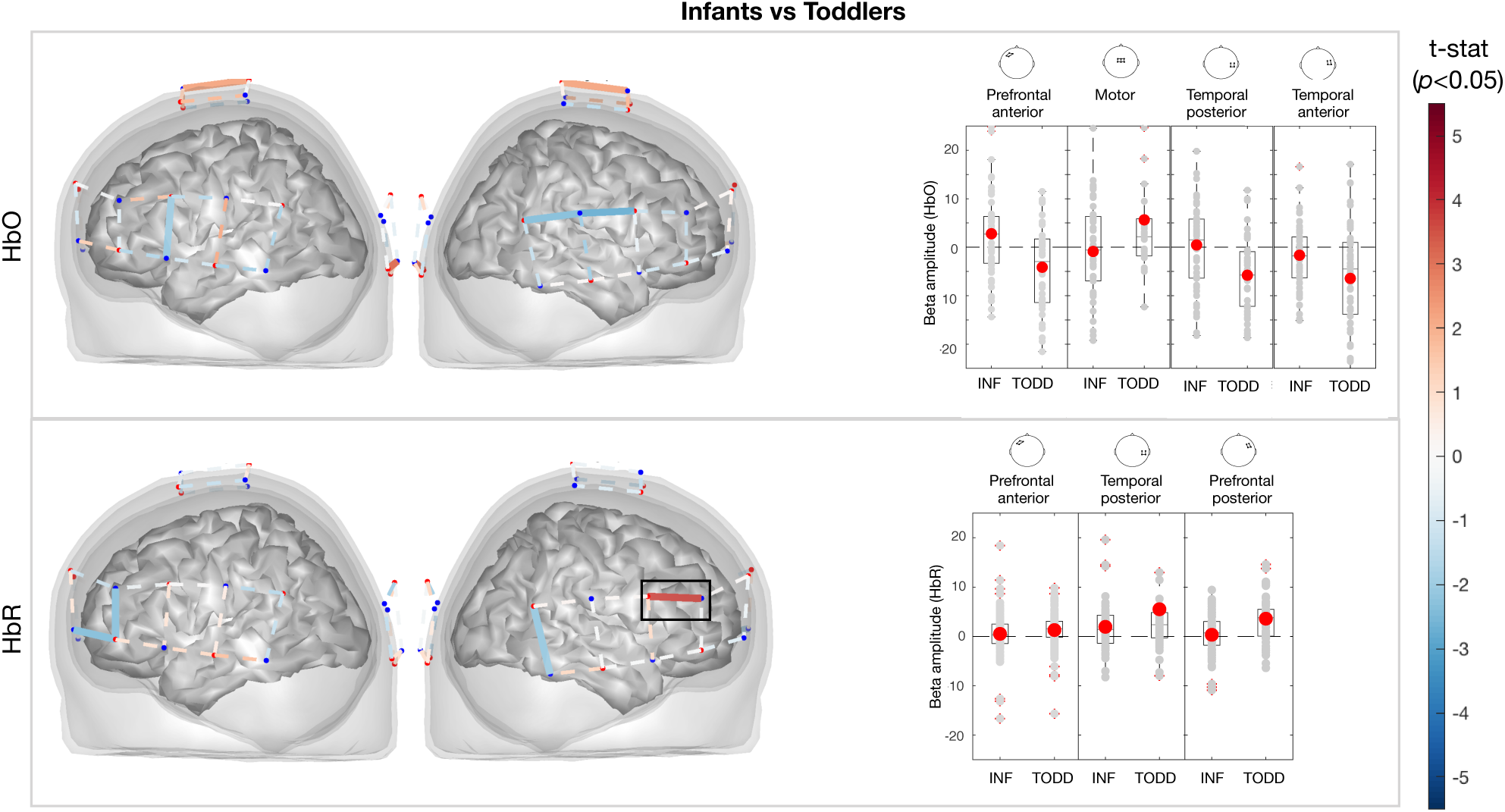
(Left) T-statistics for channels showing a significant main effect of Group for HbO (top) and HbR (bottom). The black rectangle indicates the channel that survived correction for multiple comparisons using the Benjamini–Hochberg false discovery rate (FDR; q < .05). (Right) Beta values for the significant channels (p < .05).

#### 3.2.2. Vocoded vs. Clear Condition: Main effect of condition and interaction with Age

The contrast between the vocoded and clear condition was associated with an increase in HbO activity in one channel located at the left posterior prefrontal cortex and in one channel at the left anterior temporal cortex (around left IFG, as seen in Figure 5). In addition, a decrease in HbR was observed in two channels around the left IFG, which in combination with an increase in HbO suggests an increase in brain activity in left IFG. Also, there was an increase in HbO activity in one channel on the right posterior prefrontal cortex (around right IFG), one channel on the right anterior prefrontal cortex, and in two channels at the occipital cortex (however, there was an increase in HbR activity as well). Inversely, a decrease in HbO activity was seen in the vocoded condition compared to the clear condition in one channel on the left anterior prefrontal cortex, in one channel on the right posterior temporal cortex (a decrease in HbR), and in two channels on the motor cortex.

**Figure 5.**
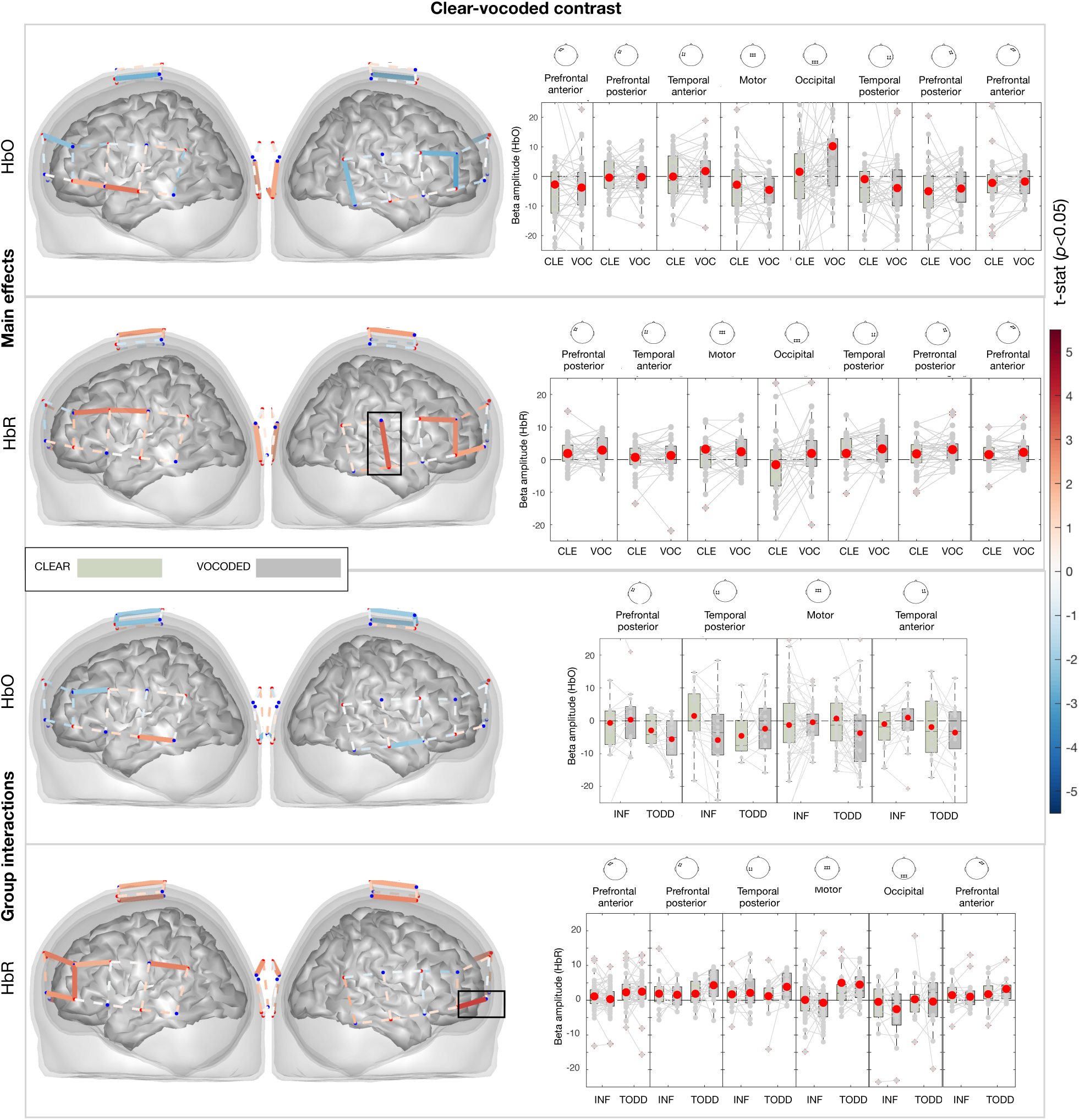
(Left) T-statistics for channels showing a significant main effect of Condition (Clear vs. Vocoded contrast) and the Condition × Group interaction, for HbO (top) and HbR (bottom). The black rectangle indicates the channel that survived correction for multiple comparisons using the Benjamini–Hochberg false discovery rate (FDR; q < .05). (Right) Corresponding beta values for channels reaching significance (p < .05).

**Figure 6.**
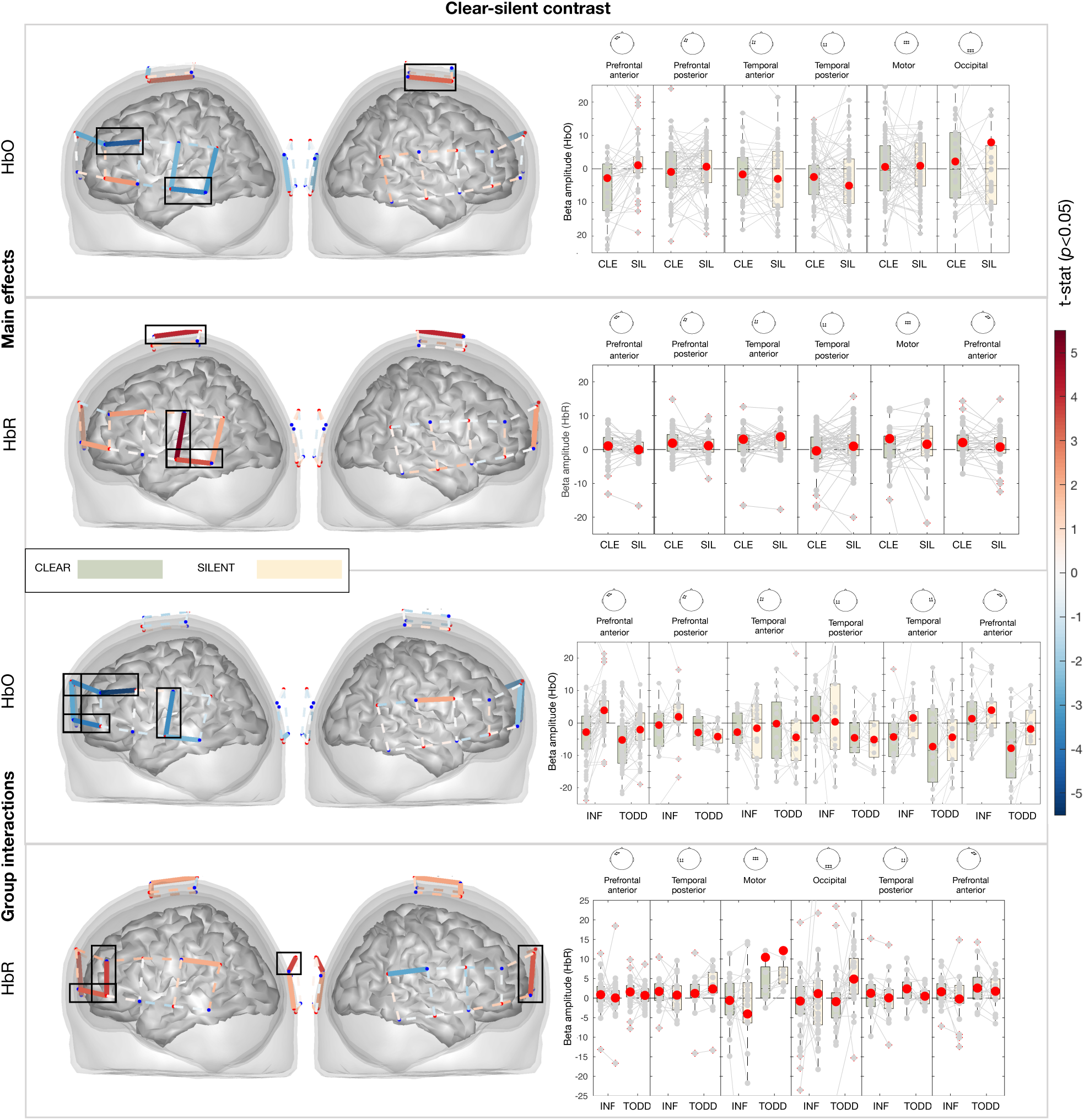
(Left) T-statistics for channels showing a significant main effect of Condition (Clear vs. Silent contrast) and the Condition × Group interaction, for HbO (top) and HbR (bottom). The black rectangle indicates the channel that survived correction for multiple comparisons using the Benjamini–Hochberg false discovery rate (FDR; q < .05). (Right) Corresponding beta values for channels reaching significance (p < .05).

None of the channels that showed a main effect for the contrast comparing vocoded to clear condition showed interactions with the group factor. However, in one channel on the left posterior prefrontal cortex, on the motor cortex and on the anterior part of the right temporal cortex, HbO activity increased for infants while it decreased for toddlers. In addition, in one channel located at the left posterior temporal cortex, HbO activity increased for toddlers but decreased for infants. Note that from the reported results, only one channel at the right temporal cortex and right prefrontal cortex passed the FDR correction for multiple comparisons for HbR (highlighted inside black boxes in Figure 5), suggesting caution with the interpretation of the results. The statistical values for each channel for each contrast are presented in tables in Supplementary Material 6.

#### 3.2.3. Silent vs. Clear condition: Main effect of condition and interaction with group

A decrease in HbO activity was seen in the silent compared to the clear condition in three channels located on the left anterior and posterior temporal cortex, and on the motor cortex (also coinciding with an increase in HbR). However, this main effect was qualified by an interaction with group, showing that toddlers were driving this decrease.

## 4 Discussion

This study examined the developmental trajectory of young children’s attentional allocation to a speaking face at varying levels of speech intelligibility and the brain activity supporting this behaviour. Our eye-tracking results showed that both infants and toddlers increased visual attention to the speaker’s mouth when hearing vocoded compared to clear speech. At the brain level, however, processing vocoded speech was supported by activity that differed by age: infants showed greater recruitment of prefrontal regions, whereas toddlers showed stronger engagement of posterior temporal regions. Finally, we observed that silent speech elicited brain responses that were distinct from those observed during vocoded audiovisual speech. These behavioural and neural response patterns are discussed in detail as follows.

### 4.1. Behavioural and neural processing of low-intelligibility speech

We predicted that a decrease in speech intelligibility would be associated with increases in visual attention to the speaker’s mouth. Our results align with our first hypothesis that this effect is independent of children’s age and level of language experience. These findings extend prior work in adults (Chandrasekaran et al., 2009) and toddlers (Król, 2018), indicating that as early as 8-months of age, looking to the mouth during acoustically degraded speech is a useful strategy to recover the missing auditory information.

At the brain level, we predicted that reduced speech intelligibility would elicit increased activity in regions associated with attentional control and effortful listening, particularly the prefrontal cortex (Davis & Johnsrude, 2003; Mushtaq et al., 2021; Peelle, 2018a), alongside reduced engagement of temporal regions surrounding the primary auditory cortex, as reported in older children under degraded listening conditions (Mushtaq et al., 2021). We further predicted that these effects may be modulated by children’s age, such that toddlers would show greater recruitment of brain regions supporting audiovisual integration, due to their greater language abilities and greater experience with audiovisual correspondences. Our results support this prediction. Although auditory degradation increased activation in brain regions associated with effortful listening and attentional control overall, the specific activation pattern differed by age. Infants showed stronger brain activation in the prefrontal cortex (left IFG and anterior prefrontal cortex) during vocoded compared to clear speech conditions, suggesting a greater reliance on domain-general attentional processes and effortful listening (Davis & Johnsrude, 2003; Giraud, 2004).

In contrast, toddlers showed stronger brain activation in the posterior temporal cortex during vocoded speech than clear speech. The posterior temporal cortex has been consistently implicated in audiovisual speech integration (Beauchamp, 2016b). Thus, one interpretation of the age-related differences observed here is that toddlers’ greater recruitment of the posterior temporal cortex reflects increased experience with audiovisual speech correspondences, enabling more efficient use of visual articulatory cues to support speech comprehension under degraded listening conditions (Campbell, 2008; Lalonde & Werner, 2021). In contrast, infants’ stronger prefrontal engagement may reflect greater processing effort and less efficient audiovisual integration, potentially due to developing associations between visual speech movements and their corresponding auditory signals. This interpretation aligns with prior developmental work showing that audiovisual speech benefits continue to mature well beyond infancy and are not adult-like until adolescence (Lalonde & Werner, 2021), suggesting that extensive experience with audiovisual speech is required before visual cues can be used optimally to compensate for degraded auditory input.

In addition to our a-priori defined regions of interest, we conducted exploratory analyses of occipital and motor cortices, as these regions have been proposed to be implicated in audiovisual speech processing in adults (Peelle, 2018), but remain largely unexplored in infants. These analyses revealed age-dependent modulation in motor regions, with toddlers showing decreased hemodynamic activity in the motor cortex during vocoded compared to clear speech. This finding differs from previous work by Kuhl et al. (2014), which reported increased motor cortex activation in 11-month-old infants when listening to unfamiliar compared to familiar languages. One possible explanation for this difference is that processing unfamiliar language and processing acoustically degraded speech rely on partially distinct mechanisms. Whereas unfamiliar language may engage motor systems involved in forming new articulatory representations, vocoded speech preserves the phonological structure of the native language and may therefore rely on existing motor representations rather than recruiting additional motor resources.

### 4.2. Behavioural and brain processing of visual-only speech

In addition to the clear and vocoded audiovisual conditions, children were also presented with a visual-only (silent speech) condition. This allowed us to characterize the behavioural and neural signatures elicited by visual speech cues alone. While this condition was not central to our design and its analyses were largely exploratory, the observed patterns could inform possible strategies used by individuals with hearing loss for whom the visible part of speech is the most reliable cue for oral speech comprehension (Weikum et al., 2007).

Children did not increase visual attention to the speaker’s mouth in the silent condition relative to clear speech. This pattern is consistent with previous findings in infants (Tan et al., 2022), but contrasts with findings from adults, who typically show increased mouth-looking during silent speech. The absence of this effect suggests that visual speech alone does not elicit the same compensatory attentional strategy as vocoded speech in early development, and that more extensive experience with audiovisual speech may be required for adult-like visual attention strategies to emerge (Lalonde & Werner, 2021). At the brain level, on the other hand, silent speech elicited age-dependent responses in speech processing brain areas. Brain activity in the prefrontal cortex (left IFG) increased during silent speech, an effect driven primarily by infants. By contrast, in the left temporal cortex, toddlers showed reduced activation during silent relative to clear speech, whereas no such modulation was observed in infants. This pattern may be in line with the findings seen in the vocoded condition, whereby infants show a greater reliance on attention-driven, effortful listening related brain areas when auditory input is entirely absent.

Across both age groups, silent speech was associated with increased activation in the motor cortex. This finding is broadly consistent with accounts of speech perception that emphasize the involvement of motor representations in processing visible speech (Bernstein & Liebenthal, 2014). However, this result should be interpreted cautiously, as increased motor activity may also have been influenced by participants’ own vocalizations during the silent trials. In this study, we did not include objective measures of children’s behaviour during the task, but this can be informative for future studies using silent stimuli with these age groups. In addition, increased activation in the occipital cortex during silent speech likely reflects enhanced visual processing reliance when auditory input is unavailable (Campbell, 2008; Emberson et al., 2015). Interestingly, silent speech did not recruit the right-hemisphere, in contrast to the pattern observed for vocoded speech, suggesting that right-hemisphere engagement may be more closely related to effortful listening under reduced intelligibility than to visual speech processing *per se*, consistent with previous results from adults (Bourguignon et al., 2020).

We note that caution is required when interpreting the developmental effects observed in our neural measures. Direct comparisons of neural responses by infants and older toddlers can be affected by developmental changes in hemodynamic responses, with infants showing predominantly positive HbO amplitudes, and toddlers displaying more negative responses. Such negative signals have been linked to mechanisms like blood stealing (McKiernan et al., 2003), inhibitory activity (Mullinger et al., 2014), or habituation (Lee, Mao, Wunderlich, et al., 2023), and may be especially relevant given toddlers’ potentially faster habituation to repeated speech stimuli. Furthermore, anatomical and physiological differences between these age groups may complicate spatial comparisons across fNIRS channels. To address this, we used region-of-interest (ROI) analyses and short video-camera recordings during calibration rather than relying on precise anatomical mapping. Future work using individual head localization or video-based channel registration could enhance spatial precision. Despite these limitations, the present design provides a valid and developmentally grounded approach to understanding how young children audiovisual speech when its intelligibility is reduced.

## 5 Conclusion

Taken together, these findings show that although infants and toddlers deploy similar behavioural strategies when auditory speech is degraded - increased attention to the speaker’s mouth - the brain mechanisms supporting this strategy differ as a function of development. Early in development, already at 8 months of age, degraded and absent auditory input is primarily associated with increased recruitment of prefrontal, domain-general attentional systems, whereas with increasing experience with language and audiovisual correspondences, speech processing is more strongly supported by posterior temporal regions implicated in audiovisual integration. Together, these findings highlight a developmental shift in speech processing when presented under challenging listening conditions. Furthermore, a greater understanding of the neural mechanisms underlying silent and vocoded speech processing can inform how these mechanisms develop in children who are affected by severe hearing loss, and those who have received cochlear implants.

## Funding info

This research was supported by the Basque Government through the BERC 2022-2025 program and Funded by the Spanish State Research Agency through BCBL Severo Ochoa excellence accreditation CEX2020-001010/AEI/10.13039/501100011033. In addition, IAS received support from the Basque Government Postdoctoral Grant (PRE_2019_1_038, and BB from the Medical Research Council Programme Grant (MR/T003057/1) and a UKRI Future Leaders fellowship (MR/S018425/1). We sincerely thank all the infants, toddlers and their families who participated in this study. We are also grateful to Elena Aguirrebengoa and Patricia Jimenez for their invaluable assistance with participant recruitment and data collection. Additionally, we thank Luca Pollonini and the Human Hearing Laboratory at the Bionics Institute, Melbourne, Australia, for their expert guidance and support in data processing.

## Author contributions

I.A.S.: Conceptualization, Methodology, Software, Investigation, Formal analysis, Visualization, Writing – original draft, Writing – review & editing. B.B.: Methodology, Software, Formal analysis, Writing – review & editing. C.C.G: Conceptualization, Methodology, Software, Formal analysis; M.C: Conceptualization, Writing – review & editing; M.K.: Supervision, Conceptualization, Methodology, Investigation, Formal analysis, Writing – review & editing.

