## Supplementary materials for "Young children rely on visual information to process degraded speech: Evidence from behavioural and neuroimaging measures"

**Supplementary Material 1**

This is the content of each of the three stories in Basque and their translated version in English.

**STORY 1:**
[Translated to English from Basque]
Good morning!
Will you help me clean the house?
We will prepare our lunch. We are hungry! Then we will buy some flowers. They are very beautiful! Do you like them? Let's put them on the table!

**STORY 2:**
[Translated to English from Basque]
Good afternoon!
We have some free time, right?
The weekend is about to start. Let's dance with our friends! We will listen to some music. The music is amazing. Do you like it?
And then we will have some chocolate!

**STORY 3:**

[Translated to English from Basque]
Good afternoon!
We will get the games ready to play, okay?
I will bring the ball. The ball is colourful!
We will run around. You are very fast! Do you want to jump? Then we will play hide-and-seek!

#### Supplementary Material 2

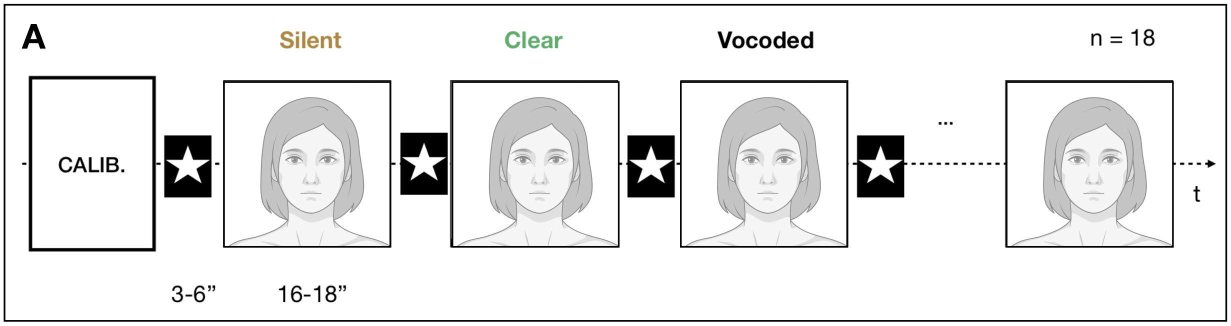


**C**

3-6” 16-18”

Version 1 (triplet ABC): **Silent** A, **Clear** B, **Vocoded** C, **Clear** A, **Vocoded** B, **Silent** C, **Vocoded** A, **Silent** B, **Clear** C [all x2]

Version 2 (triplet ABC): **Silent** A, **Clear** B, **Vocoded** C, **Clear** A, **Silent** B, **Vocoded** C, **Clear** A, **Vocoded** B, **Silent** C [all x2]

Version 3 (triplet BCA): **Silent** B, **Clear** C, **Vocoded** A, **Clear** B, **Vocoded** C, **Silent** A, **Vocoded** B, **Silent** C, **Clear** A [all x2]

Version 4 (triplet BCA): **Silent** B, **Clear** C, **Vocoded** A, **Clear** B, **Silent** C, **Vocoded** A, **Clear** B, **Vocoded** C, **Silent** A [all x2]

Version 5 (triplet CAB): **Silent** C, **Clear** A, **Vocoded** B, **Clear** C, **Vocoded** A, **Silent** B, **Vocoded** C, **Silent** A, **Clear** B [all x2]

Version 6 (triplet CAB): **Silent** C, **Clear** A, **Vocoded** B, **Clear** C, **Silent** A, **Vocoded** B, **Clear** C, **Vocoded** A, **Silent** B [all x2]

*A) Illustration of the experimental procedure, showing only the first three trials out of 18. From the 4$^{th}$ to the 18$^{th}$ trial, the conditions were ordered minimizing repeating consecutive conditions and two different orders of presentation were created, to avoid any presentation order effect. The first three trials are common for both orders of presentation: starting with a silent trial, followed by a normal trial and a vocoded trial. In addition, the order of the three monologues (different in content) was also counterbalanced to avoid effects driven by the content of the monologues. C) To minimize the influence of the content of the stories (labelled as: A, B, C), the order of the stories was presented sequentially, creating three orders in which the content was different. The combination of the 2 different condition presentation orders with the 3 different story-triplet combinations resulted in 6 different versions, that were repeated twice making a total of 18 trials, as each 9 trial sequence was repeated twice.*

**Supplementary Material 3**

Eye-tracking quality assessment:

From the included participants, infants had a total loss of trials of 48.8 %, and toddlers a total loss of trials of 66.9 %.

fNIRS montage creation and quality assessment:

To analyse the fNIRS signals, the channels were grouped into regions of interest (ROI) using the devFOLD application in Matlab (Fu & Richards, 2021) and the AAL brain atlas, choosing a specificity of 30% (default). For the temporal cortex ROI, we placed channels over the middle and superior temporal cortex (seen as pink and purple dots in Figure 2B). For the prefrontal cortex ROI, we placed channels over the inferior frontal gyrus (pars triangularis) (seen as orange and yellow dots in Figure 2B). For the motor cortex ROI, we placed channels over the right and left supplementary motor cortex (seen as white dots in Figure 2B). For the occipital cortex ROI, we placed channels that corresponded to the middle and superior occipital cortex (seen as black dots in Figure 2B).

The following two quality assessments were performed to the fNIRS data, using *nirs-toolbox* (Santosa et al., 2018) in Matlab R2018a: 1) At the channel level: Channel quality was assessed using the scalp coupling index (SCI), computed first by isolating the cardiac oscillations bandpassing the two wavelength signals between 0.5 and 2.5 Hz and then as the zero-lag cross-correlation between the two wavelength signals. If the SCI was above 0.7, the channel was included in the analysis (Pollonini et al., 2016). 2) At the trial level: The SCI-based channel quality assessment was unreliable for the youngest participants because their higher heart rates (~2 Hz) exceeded the detectable frequency range given the 3.9 Hz sampling rate. As a result, cardiac oscillations could not be reliably captured, making SCI an unsuitable quality marker for these subjects (see Supplementary material 13 for more details on the calculation of the heart-beat information). To overcome this limitation, on top of using the SCI measure on all participants (which was not really informative for the youngest subjects), in addition to the SCI, the heartbeat information was also used for the inclusion of trials. If the cardiac signal was present in at least $1/3^{rd}$ of the trial, the trial was included. If a trial was not included, it was converted into *nan* values (Lee et al., 2023). In addition, the trigger corresponding to that trial (epoch) was deleted – therefore not used for the GLM analyses.

Participants were included only if they had good eye tracking quality, if they had more than 50% of good quality channels, if they had at least 2 good channels in the temporal ROI (as a control measure for linguistic processing) and if they had at least two good trials per condition (see Supplementary materials 2 for individual-level quality assessment). After applying the exclusion criteria, the final sample for the fNIRS analysis consisted of 18 participants (out of 23 with good ET quality) in the infant group, as one participant had to be excluded due to the wrong selection of montage, and 4 did not have at least two good epochs per condition. 18 participants (out of 20 with good ET quality) were included in the toddler group, as one participant did not have at least two good epochs for each condition, and one participant did not have at least two good channels in the temporal ROI. From the included participants, the infant group had a total loss of channels of 0.6 %, and the toddler group had a total loss of channels of 16.3 %. From the included participants, the infant group had a 28% of trial loss and toddlers a 12.3% of trial loss.

### Supplementary Material 4

| Participant | ET | | | NIRS | | | | | | | | | |
|---|---|---|---|---|---|---|---|---|---|---|---|---|---|
| | | | | Epoch based quality ass. | | | SCI based quality ass. | | | | | | |
| | AV | AVdeg | V | AV (6) | AV degr (6) | V (6) | SCI all chann | Left PFC (6) | Right PFC (6) | eft temp (7) | ght temp (7) | Motor (7) | Occi (7) |
| BB03 | 5 | 5 | 3 | 3 | 2 | 2 | 0 | 0 | 0 | 0 | 0 | 0 | 0 |
| BB04 | 4 | 3 | 1 | 4 | 1 | 1 | 0 | 0 | 0 | 0 | 0 | 0 | 0 |
| BB05 | 6 | 6 | 4 | 2 | 2 | 2 | 0 | 0 | 0 | 0 | 0 | 0 | 0 |
| BB06 | 5 | 4 | 4 | 3 | 1 | 2 | 0 | 0 | 0 | 0 | 0 | 0 | 0 |
| BB07 | 2 | 1 | 1 | 3 | 1 | 2 | 2 | 0 | 0 | 0 | 1 | 0 | 1 |
| BB09 | 2 | 0 | 1 | Wrong montage | | | | | | | | | |
| BB11 | 2 | 5 | 4 | 1 | 0 | 0 | 0 | 0 | 0 | 0 | 0 | 0 | 0 |
| BB13 | 1 | 1 | 2 | 0 | 1 | 0 | 0 | 0 | 0 | 0 | 0 | 0 | 0 |
| BB14 | 2 | 3 | 2 | 0 | 1 | 0 | 0 | 0 | 0 | 0 | 0 | 0 | 0 |
| BB15 | 4 | 4 | 3 | 0 | 1 | 0 | 0 | 0 | 0 | 0 | 0 | 0 | 0 |
| BB16 | 1 | 2 | 1 | 5 | 5 | 5 | 0 | 0 | 0 | 0 | 0 | 0 | 0 |
| BB17 | 4 | 3 | 3 | 1 | 1 | 0 | 0 | 0 | 0 | 0 | 0 | 0 | 0 |
| BB19 | 4 | 5 | 4 | 3 | 3 | 3 | 0 | 0 | 0 | 0 | 0 | 0 | 0 |
| BB21 | 1 | 1 | 2 | 4 | 4 | 3 | 0 | 0 | 0 | 0 | 0 | 0 | 0 |
| BB22 | 5 | 3 | 4 | 3 | 0 | 1 | 0 | 0 | 0 | 0 | 0 | 0 | 0 |
| BB23 | 2 | 2 | 1 | 5 | 3 | 3 | 0 | 0 | 0 | 0 | 0 | 0 | 0 |
| BB24_2 | 5 | 4 | 4 | 0 | 0 | 0 | 0 | 0 | 0 | 0 | 0 | 0 | 2 |
| BB27 | 2 | 4 | 3 | 1 | 2 | 1 | 0 | 0 | 0 | 0 | 0 | 0 | 0 |
| BB29 | 5 | 3 | 1 | 6 | 5 | 5 | 0 | 0 | 0 | 0 | 1 | 0 | 1 |
| BB30 | 5 | 4 | 6 | 2 | 3 | 2 | 0 | 0 | 0 | 0 | 0 | 0 | 0 |
| BB31 | 0 | 1 | 3 | 6 | 6 | 6 | 0 | 0 | 0 | 0 | 0 | 0 | 0 |
| BB32 | 2 | 2 | 1 | 2 | 4 | 2 | 0 | 0 | 0 | 0 | 0 | 0 | 0 |
| BB33 | 3 | 3 | 3 | 2 | 2 | 2 | 3 | 0 | 0 | 0 | 0 | 2 | 1 |
| BBOLD01 | 5 | 6 | 5 | 1 | 1 | 0 | 9 | 0 | 1 | 2 | 1 | 3 | 2 |
| BBOLD03 | 4 | 4 | 2 | 1 | 0 | 0 | 6 | 2 | 0 | 0 | 3 | 0 | 1 |
| BBOLD04 | 4 | 4 | 4 | 1 | 0 | 0 | 0 | 0 | 0 | 0 | 0 | 0 | 0 |
| BBOLD05 | 5 | 6 | 6 | 0 | 0 | 0 | 16 | 0 | 2 | 4 | 6 | 2 | 2 |
| BBOLD07 | 6 | 6 | 1 | 0 | 0 | 0 | 4 | 0 | 0 | 0 | 0 | 0 | 4 |
| BBOLD09 | 5 | 5 | 4 | 2 | 0 | 0 | 7 | 0 | 0 | 0 | 3 | 2 | 2 |
| BBOLD10 | 6 | 6 | 4 | 0 | 0 | 0 | 16 | 2 | 0 | 6 | 3 | 0 | 5 |
| BBOLD12 | 4 | 4 | 5 | 1 | 0 | 1 | 4 | 0 | 0 | 0 | 0 | 0 | 4 |
| BBOLD13 | 5 | 4 | 4 | 0 | 0 | 0 | 3 | 0 | 0 | 0 | 0 | 3 | 0 |
| BBOLD15 | 6 | 6 | 6 | 0 | 0 | 0 | 8 | 0 | 0 | 0 | 2 | 4 | 2 |
| BBOLD16 | 3 | 4 | 4 | 2 | 3 | 3 | 6 | 0 | 0 | 1 | 2 | 1 | 2 |
| BBOLD17 | 3 | 2 | 1 | 2 | 3 | 3 | 20 | 1 | 3 | 5 | 5 | 3 | 3 |
| BBOLD18 | 5 | 6 | 5 | 1 | 0 | 0 | 1 | 0 | 0 | 0 | 0 | 0 | 1 |
| BBOLD20 | 2 | 2 | 3 | 6 | 3 | 5 | 15 | 0 | 1 | 5 | 2 | 4 | 3 |
| BBOLD22 | 6 | 6 | 4 | 1 | 0 | 0 | 4 | 0 | 0 | 2 | 1 | 0 | 1 |
| BBOLD26 | 1 | 1 | 2 | 1 | | 0 | 8 | 0 | 0 | 0 | 1 | 2 | 5 |
| BBOLD30 | 3 | 2 | 2 | 0 | 0 | 1 | 3 | 0 | 0 | 2 | 0 | 1 | 0 |
| BBOLD31 | 2 | 3 | 2 | 2 | 0 | 0 | 13 | 0 | 0 | 1 | 1 | 5 | 6 |
| BBOLD32 | 3 | 4 | 2 | 2 | 0 | 0 | 17 | 2 | 2 | 3 | 1 | 5 | 4 |
| BBOLD33 | 5 | 6 | 5 | 0 | 0 | 0 | 9 | 0 | 0 | 0 | 0 | 3 | 6 |

*Figure 1. Included participants for ET analysis, representing how many GOOD trials were included per condition (2 good trials per at least one condition was the criteria for a participant to be included). From those participants, the rows in red represent those participants that did not pass the NIRS quality criteria, either due to having too many BAD epochs (>4 out of 6) or having too many BAD channels in the temporal ROI (>40) (coincided that all participants with good ET also had >50% good quality NIRS channels)*

**Supplementary Material 5**

**Calculation of the heart-beat information when sampling frequency lower (3.9 Hz) than that needed for the high frequency heart-beat of youngest participants.**

The low sampling frequency (~3.9 Hz) of our fNIRS recordings only permits a resolution of 1.95 Hz (half the sampling frequency, due to the Nyquist theorem). Given that during infancy the heart beats at faster rates, sometimes higher than 1.95 Hz (Fleming et al., 2011), the quality assessment used by Scalp Coupling Index (which uses heart beat information to assess fNIRS signal quality), is not always effective. Therefore, the heart beat signal was reconstructed using the whole-brain heartbeat information, as a measure of quality of the recording over time, following previous work by (Lee et al., 2023).

The illumination pattern of the NIRScout machine lights each source sequentially, one source at a time. This sequential illumination pattern was used to increase the number of points of the recording. Instead of grouping all source lightening times at once (the default outcome given by the NIRScout machine), each point in time was represented when it was recorded. This provided many more points in time, and as the cardiac oscillation is a signal that is present across the cortex (regardless of the location in the cortex), it could be reconstructed with a higher frequency resolution. Finally, this information was used to assess trial-by-trial quality.

**Supplementary Material 6**

At the group level, statistical analyses were conducted using a linear mixed-effects model specified as:

$$Amplitude(\beta) \sim Condition\ (clear, vocoded, silent) \times Group\ (infants, toddlers) + (1 \mid Subject)$$

where condition and group were entered as fixed effects, including their interaction, and subject was included as a random intercept.

For each channel, parameter estimates are reported for all contrasts. Channels reaching statistical significance for a given contrast are highlighted in light yellow.

The contrasts are defined as follows:

- b_AVD: contrast between the clear and vocoded conditions
- c_V: contrast between the clear and silent conditions
- group: main effect of group (infants vs. toddlers)
- b_AVD × group: interaction between group and the clear vs. vocoded contrast
- c_V × group: interaction between group and the clear vs. silent contrast

*Channel statistics, output from the glm model by block .*

| Source | Detector | Signal type | Contrast | Beta | T-stat | P-value | Q-value |
|---|---|---|---|---|---|---|---|
| 1 | 1 | 'hbo' | 'b_AVD' | -0.528037 | -0.8392264 | 0.40328589 | 0.70655958 |
| 1 | 1 | 'hbo' | 'c_V' | -0.0308571 | -0.048669 | 0.96127729 | 0.9808952 |
| 1 | 1 | 'hbo' | 'group' | -1.231544 | -0.9812851 | 0.32875152 | 0.63596069 |
| 1 | 1 | 'hbo' | 'b_AVD:grou | -1.7205902 | -1.0965639 | 0.27538924 | 0.59229589 |
| 1 | 1 | 'hbo' | 'c_V:group' | -4.474887 | -2.8422557 | **0.00540156** | 0.08199133 |
| 1 | 1 | 'hbr' | 'b_AVD' | -0.5301077 | -1.7025089 | 0.09167618 | 0.35175888 |
| 1 | 1 | 'hbr' | 'c_V' | 0.49380521 | 1.57935982 | 0.1173198 | 0.39501289 |
| 1 | 1 | 'hbr' | 'group' | 0.48856448 | 0.73134033 | 0.46623187 | 0.72777559 |
| 1 | 1 | 'hbr' | 'b_AVD:grou | 1.28888301 | 1.73859714 | 0.08509375 | 0.34536956 |
| 1 | 1 | 'hbr' | 'c_V:group' | 1.15065104 | 1.54749405 | 0.12481132 | 0.40589047 |
| 1 | 2 | 'hbo' | 'b_AVD' | -1.4979907 | -2.2965486 | **0.0236688** | 0.19445821 |
| 1 | 2 | 'hbo' | 'c_V' | -1.9032768 | -2.8338829 | **0.00553441** | 0.08199133 |
| 1 | 2 | 'hbo' | 'group' | -0.3317967 | -0.2545755 | 0.79955856 | 0.92971925 |
| 1 | 2 | 'hbo' | 'b_AVD:grou | -2.4796745 | -1.6191743 | 0.10846781 | 0.38738503 |
| 1 | 2 | 'hbo' | 'c_V:group' | -5.806951 | -3.6584443 | **0.00040198** | 0.01582182 |
| 1 | 2 | 'hbr' | 'b_AVD' | -0.4819376 | -1.3566806 | 0.17784879 | 0.49061735 |
| 1 | 2 | 'hbr' | 'c_V' | -0.4425178 | -1.2441578 | 0.21626535 | 0.54026705 |
| 1 | 2 | 'hbr' | 'group' | -0.8300132 | -1.1310407 | 0.2606646 | 0.58576314 |
| 1 | 2 | 'hbr' | 'b_AVD:grou | 2.42946988 | 2.81192145 | **0.00589724** | 0.0820438 |
| 1 | 2 | 'hbr' | 'c_V:group' | 1.69418065 | 1.95903448 | 0.05281214 | 0.27083146 |
| 2 | 1 | 'hbo' | 'b_AVD' | 0.13664255 | 0.23620226 | 0.81374479 | 0.93266399 |
| 2 | 1 | 'hbo' | 'c_V' | 0.44709532 | 0.76744654 | 0.44457177 | 0.71417152 |
| 2 | 1 | 'hbo' | 'group' | 1.55883394 | 1.35724867 | 0.17766885 | 0.49061735 |
| 2 | 1 | 'hbo' | 'b_AVD:grou | -2.1543107 | -1.4608989 | 0.14708706 | 0.44536457 |
| 2 | 1 | 'hbo' | 'c_V:group' | -5.7003953 | -3.8555235 | **0.00020128** | 0.01582182 |
| 2 | 1 | 'hbr' | 'b_AVD' | 0.34028397 | 1.09649881 | 0.27541759 | 0.59229589 |
| 2 | 1 | 'hbr' | 'c_V' | 0.705074 | 2.25536635 | **0.02622511** | 0.20173161 |
| 2 | 1 | 'hbr' | 'group' | -1.4616892 | -2.2928462 | **0.0238893** | 0.19445821 |
| 2 | 1 | 'hbr' | 'b_AVD:grou | 1.86583985 | 2.45502196 | **0.01576272** | 0.16166889 |
| 2 | 1 | 'hbr' | 'c_V:group' | 2.48850251 | 3.25008183 | **0.00155905** | 0.04157456 |
| 2 | 2 | 'hbo' | 'b_AVD' | -0.7237414 | -1.300993 | 0.19616377 | 0.51963913 |
| 2 | 2 | 'hbo' | 'c_V' | 0.66042695 | 1.17458065 | 0.24287095 | 0.55832403 |
| 2 | 2 | 'hbo' | 'group' | -0.7950342 | -0.7164835 | 0.4753143 | 0.72777559 |
| 2 | 2 | 'hbo' | 'b_AVD:grou | -1.9980734 | -1.3976496 | 0.16522137 | 0.47545718 |
| 2 | 2 | 'hbo' | 'c_V:group' | -1.4833346 | -1.0312262 | 0.30485015 | 0.60586223 |
| 2 | 2 | 'hbr' | 'b_AVD' | 0.40790749 | 1.57075749 | 0.11930598 | 0.39768662 |
| 2 | 2 | 'hbr' | 'c_V' | 0.29168302 | 1.10779194 | 0.27053192 | 0.58842528 |
| 2 | 2 | 'hbr' | 'group' | -1.1715611 | -2.1535409 | **0.03360924** | 0.22785926 |
| 2 | 2 | 'hbr' | 'b_AVD:grou | 1.85448447 | 2.98582707 | **0.00353369** | 0.06671597 |
| 2 | 2 | 'hbr' | 'c_V:group' | 2.32222042 | 3.69375028 | **0.00035575** | 0.01582182 |
| 2 | 3 | 'hbo' | 'b_AVD' | 1.21726334 | 2.01573432 | **0.04643198** | 0.25090528 |
| 2 | 3 | 'hbo' | 'c_V' | 1.4656543 | 2.38691977 | **0.01881468** | 0.17502023 |
| 2 | 3 | 'hbo' | 'group' | -1.9268989 | -1.5961983 | 0.11350826 | 0.39140779 |
| 2 | 3 | 'hbo' | 'b_AVD:grou | 0.98908509 | 0.63703064 | 0.52551826 | 0.74541597 |
| 2 | 3 | 'hbo' | 'c_V:group' | -1.116899 | -0.7113763 | 0.47845907 | 0.72777559 |
| 2 | 3 | 'hbr' | 'b_AVD' | 0.39629968 | 1.12338421 | 0.26388592 | 0.58765872 |
| 2 | 3 | 'hbr' | 'c_V' | 0.08415121 | 0.23619639 | 0.81374934 | 0.93266399 |
| 2 | 3 | 'hbr' | 'group' | 0.56597894 | 0.77840021 | 0.43811764 | 0.71417152 |
| 2 | 3 | 'hbr' | 'b_AVD:grou | 0.02584339 | 0.0296412 | 0.97641056 | 0.98893271 |
| 2 | 3 | 'hbr' | 'c_V:group' | 0.32166513 | 0.366028 | 0.7150949 | 0.87399027 |

| Source | Detector | Signal type | Contrast | Beta | T-stat | P-value | Q-value |
|---|---|---|---|---|---|---|---|
| 3 | 2 | 'hbo' | 'b_AVD' | -1.0098491 | -1.8410595 | 0.06849106 | 0.311323 |
| 3 | 2 | 'hbo' | 'c_V' | -2.6248358 | -4.8465447 | **4.46E-06** | 0.00059405 |
| 3 | 2 | 'hbo' | 'group' | 1.67734928 | 1.54850306 | 0.12456844 | 0.40589047 |
| 3 | 2 | 'hbo' | 'b_AVD:grou | -2.8361247 | -2.116144 | **0.03674263** | 0.24093531 |
| 3 | 2 | 'hbo' | 'c_V:group' | -7.1654411 | -5.4084432 | **4.15E-07** | 0.00016583 |
| 3 | 2 | 'hbr' | 'b_AVD' | 0.97417967 | 2.95725488 | **0.00384949** | 0.06694774 |
| 3 | 2 | 'hbr' | 'c_V' | 0.79795568 | 2.388891 | **0.01871944** | 0.17502023 |
| 3 | 2 | 'hbr' | 'group' | -0.27924 | -0.4205364 | 0.67496974 | 0.85439208 |
| 3 | 2 | 'hbr' | 'b_AVD:grou | 1.63939598 | 2.03391053 | **0.04453129** | 0.25090528 |
| 3 | 2 | 'hbr' | 'c_V:group' | 1.40737561 | 1.72723244 | 0.08712349 | 0.34536956 |
| 3 | 3 | 'hbo' | 'b_AVD' | 0.68399568 | 0.81093539 | 0.41927265 | 0.71004527 |
| 3 | 3 | 'hbo' | 'c_V' | -1.038304 | -1.215176 | 0.22707714 | 0.54840223 |
| 3 | 3 | 'hbo' | 'group' | -3.3712433 | -2.01001 | **0.04704474** | 0.25090528 |
| 3 | 3 | 'hbo' | 'b_AVD:grou | 0.81979298 | 0.3793358 | 0.70521944 | 0.87367661 |
| 3 | 3 | 'hbo' | 'c_V:group' | -0.5345941 | -0.2446419 | 0.80722043 | 0.93266399 |
| 3 | 3 | 'hbr' | 'b_AVD' | 0.38884743 | 1.06075657 | 0.29128156 | 0.59258348 |
| 3 | 3 | 'hbr' | 'c_V' | 0.08418664 | 0.22454017 | 0.82278185 | 0.93518807 |
| 3 | 3 | 'hbr' | 'group' | 0.60611764 | 0.80534313 | 0.42247694 | 0.71004527 |
| 3 | 3 | 'hbr' | 'b_AVD:grou | -0.3012873 | -0.3475829 | 0.72886273 | 0.87613134 |
| 3 | 3 | 'hbr' | 'c_V:group' | -0.0704613 | -0.0795065 | 0.93678405 | 0.97581672 |
| 3 | 4 | 'hbo' | 'b_AVD' | -0.5850095 | -0.656936 | 0.51268723 | 0.74472351 |
| 3 | 4 | 'hbo' | 'c_V' | -0.5868506 | -0.652386 | 0.51560552 | 0.74472351 |
| 3 | 4 | 'hbo' | 'group' | -2.6037425 | -1.5108687 | 0.13388383 | 0.41838696 |
| 3 | 4 | 'hbo' | 'b_AVD:grou | 1.62963683 | 0.7247181 | 0.47026811 | 0.72777559 |
| 3 | 4 | 'hbo' | 'c_V:group' | 1.50938802 | 0.6652234 | 0.50739446 | 0.74472351 |
| 3 | 4 | 'hbr' | 'b_AVD' | 1.06004786 | 2.64172548 | **0.00953409** | 0.10997458 |
| 3 | 4 | 'hbr' | 'c_V' | 0.6781166 | 1.68779408 | 0.0944767 | 0.35318392 |
| 3 | 4 | 'hbr' | 'group' | 0.01519674 | 0.01884672 | 0.98499984 | 0.99244316 |
| 3 | 4 | 'hbr' | 'b_AVD:grou | 0.23264487 | 0.23645195 | 0.81355157 | 0.93266399 |
| 3 | 4 | 'hbr' | 'c_V:group' | 1.8312315 | 1.8493173 | 0.06727956 | 0.30933131 |
| 4 | 3 | 'hbo' | 'b_AVD' | 1.95749816 | 2.9732732 | **0.00366938** | 0.06671597 |
| 4 | 3 | 'hbo' | 'c_V' | -1.1714955 | -1.7305814 | 0.08652129 | 0.34536956 |
| 4 | 3 | 'hbo' | 'group' | -1.2538669 | -0.9635159 | 0.33754535 | 0.64250315 |
| 4 | 3 | 'hbo' | 'b_AVD:grou | 1.79768161 | 1.05950671 | 0.29184736 | 0.59258348 |
| 4 | 3 | 'hbo' | 'c_V:group' | -1.7328697 | -1.001259 | 0.31904807 | 0.62866614 |
| 4 | 3 | 'hbr' | 'b_AVD' | -0.0380401 | -0.1155039 | 0.90827065 | 0.9636272 |
| 4 | 3 | 'hbr' | 'c_V' | 0.04871063 | 0.14450027 | 0.88538781 | 0.95949752 |
| 4 | 3 | 'hbr' | 'group' | 0.63076156 | 0.93746964 | 0.35071005 | 0.64647014 |
| 4 | 3 | 'hbr' | 'b_AVD:grou | 0.28780939 | 0.3592223 | 0.72016416 | 0.87399027 |
| 4 | 3 | 'hbr' | 'c_V:group' | -1.3057431 | -1.6004202 | 0.11256828 | 0.39140779 |
| 4 | 4 | 'hbo' | 'b_AVD' | 0.44329355 | 0.60636988 | 0.54560329 | 0.7604227 |
| 4 | 4 | 'hbo' | 'c_V' | -1.8275561 | -2.4772916 | **0.01486523** | 0.15647607 |
| 4 | 4 | 'hbo' | 'group' | 2.67611536 | 1.96056968 | 0.05263006 | 0.27083146 |
| 4 | 4 | 'hbo' | 'b_AVD:grou | -1.4019067 | -0.7581081 | 0.45011737 | 0.7201878 |
| 4 | 4 | 'hbo' | 'c_V:group' | -6.4213097 | -3.4378949 | **0.00084709** | 0.02606441 |
| 4 | 4 | 'hbr' | 'b_AVD' | 0.59333431 | 1.41048766 | 0.16140869 | 0.46785127 |
| 4 | 4 | 'hbr' | 'c_V' | 2.18765916 | 5.14476262 | **1.29E-06** | 0.00025705 |
| 4 | 4 | 'hbr' | 'group' | 0.4428938 | 0.52117835 | 0.60336129 | 0.80717229 |
| 4 | 4 | 'hbr' | 'b_AVD:grou | 1.14034635 | 1.09236213 | 0.27722241 | 0.59258348 |
| 4 | 4 | 'hbr' | 'c_V:group' | 1.18381707 | 1.12205869 | 0.26444642 | 0.58765872 |

| Source | Detector | Signal type | Contrast | Beta | T-stat | P-value | Q-value |
|---|---|---|---|---|---|---|---|
| 4 | 5 | 'hbo' | 'b_AVD' | 0.79778301 | 0.88538146 | 0.37801325 | 0.68441824 |
| 4 | 5 | 'hbo' | 'c_V' | -3.3686795 | -3.6433534 | **0.00042342** | 0.01582182 |
| 4 | 5 | 'hbo' | 'group' | -3.0947453 | -1.740203 | 0.08481009 | 0.34536956 |
| 4 | 5 | 'hbo' | 'b_AVD:grou | 5.50471593 | 2.3619575 | **0.02005836** | 0.18234875 |
| 4 | 5 | 'hbo' | 'c_V:group' | -5.9064585 | -2.4844976 | **0.01458476** | 0.15647607 |
| 4 | 5 | 'hbr' | 'b_AVD' | 0.268277 | 0.63999192 | 0.52359892 | 0.74533654 |
| 4 | 5 | 'hbr' | 'c_V' | 1.49616143 | 3.48132997 | **0.00073323** | 0.02444095 |
| 4 | 5 | 'hbr' | 'group' | 1.36123762 | 1.61259442 | 0.10989259 | 0.38900031 |
| 4 | 5 | 'hbr' | 'b_AVD:grou | 1.28105638 | 1.28375472 | 0.20210826 | 0.52156971 |
| 4 | 5 | 'hbr' | 'c_V:group' | -0.5115246 | -0.4957263 | 0.62114358 | 0.8161078 |
| 5 | 4 | 'hbo' | 'b_AVD' | 0.86183893 | 1.17609613 | 0.24226761 | 0.55832403 |
| 5 | 4 | 'hbo' | 'c_V' | -1.320697 | -1.7657903 | 0.0803942 | 0.34536956 |
| 5 | 4 | 'hbo' | 'group' | 0.00159492 | 0.00109455 | 0.99912879 | 0.99912879 |
| 5 | 4 | 'hbo' | 'b_AVD:grou | 0.8679704 | 0.46364679 | 0.64387914 | 0.82814037 |
| 5 | 4 | 'hbo' | 'c_V:group' | -1.4582687 | -0.7676383 | 0.44445831 | 0.71417152 |
| 5 | 4 | 'hbr' | 'b_AVD' | 0.32184789 | 0.97409917 | 0.33228946 | 0.63596069 |
| 5 | 4 | 'hbr' | 'c_V' | 0.11861836 | 0.35804551 | 0.72104198 | 0.87399027 |
| 5 | 4 | 'hbr' | 'group' | -0.9984415 | -1.4572708 | 0.14808372 | 0.44536457 |
| 5 | 4 | 'hbr' | 'b_AVD:grou | 2.23328864 | 2.8089378 | **0.00594818** | 0.0820438 |
| 5 | 4 | 'hbr' | 'c_V:group' | 1.61425079 | 2.03101096 | **0.04482996** | 0.25090528 |
| 5 | 5 | 'hbo' | 'b_AVD' | -1.7657492 | -1.8857218 | 0.06215002 | 0.30276266 |
| 5 | 5 | 'hbo' | 'c_V' | -2.9613709 | -3.1115241 | **0.00240863** | 0.05352504 |
| 5 | 5 | 'hbo' | 'group' | -3.0028117 | -1.6226667 | 0.10771764 | 0.38738503 |
| 5 | 5 | 'hbo' | 'b_AVD:grou | -0.6744469 | -0.2840186 | 0.77696629 | 0.91662597 |
| 5 | 5 | 'hbo' | 'c_V:group' | -2.3475367 | -0.9754219 | 0.33163634 | 0.63596069 |
| 5 | 5 | 'hbr' | 'b_AVD' | 0.50394072 | 1.18363717 | 0.23928127 | 0.55646806 |
| 5 | 5 | 'hbr' | 'c_V' | 0.89961314 | 2.060988 | **0.04182349** | 0.25090528 |
| 5 | 5 | 'hbr' | 'group' | -1.0517907 | -1.2095135 | 0.22923447 | 0.54906459 |
| 5 | 5 | 'hbr' | 'b_AVD:grou | 1.30922514 | 1.23520679 | 0.21956369 | 0.54026705 |
| 5 | 5 | 'hbr' | 'c_V:group' | 1.89034513 | 1.74131481 | 0.08461416 | 0.34536956 |
| 6 | 6 | 'hbo' | 'b_AVD' | -1.8625777 | -2.5650666 | **0.01175645** | 0.13062723 |
| 6 | 6 | 'hbo' | 'c_V' | 0.69743161 | 0.95218147 | 0.34323409 | 0.64301855 |
| 6 | 6 | 'hbo' | 'group' | 2.49579767 | 1.73441516 | 0.08583609 | 0.34536956 |
| 6 | 6 | 'hbo' | 'b_AVD:grou | -4.2558855 | -2.2733046 | **0.02508327** | 0.19673154 |
| 6 | 6 | 'hbo' | 'c_V:group' | -2.8621972 | -1.5200133 | 0.13157151 | 0.41439845 |
| 6 | 6 | 'hbr' | 'b_AVD' | -0.4567613 | -1.0832037 | 0.28124725 | 0.59258348 |
| 6 | 6 | 'hbr' | 'c_V' | 0.72713453 | 1.69302889 | 0.09347254 | 0.35272658 |
| 6 | 6 | 'hbr' | 'group' | -1.0307949 | -1.2162948 | 0.22665265 | 0.54840223 |
| 6 | 6 | 'hbr' | 'b_AVD:grou | 1.02313744 | 0.95063525 | 0.34401492 | 0.64301855 |
| 6 | 6 | 'hbr' | 'c_V:group' | 1.9091761 | 1.75764335 | 0.08177918 | 0.34536956 |
| 6 | 7 | 'hbo' | 'b_AVD' | -0.5863632 | -0.7132641 | 0.47729532 | 0.72777559 |
| 6 | 7 | 'hbo' | 'c_V' | -1.8393367 | -2.210538 | **0.02927994** | 0.2116249 |
| 6 | 7 | 'hbo' | 'group' | 1.96198606 | 1.23360216 | 0.22015882 | 0.54026705 |
| 6 | 7 | 'hbo' | 'b_AVD:grou | -4.9718347 | -2.3200152 | **0.02231249** | 0.19445821 |
| 6 | 7 | 'hbo' | 'c_V:group' | -1.3889746 | -0.6434047 | 0.52139146 | 0.74484494 |
| 6 | 7 | 'hbr' | 'b_AVD' | 0.55653233 | 1.07549026 | 0.28466812 | 0.59258348 |
| 6 | 7 | 'hbr' | 'c_V' | -0.2991646 | -0.5738111 | 0.56734651 | 0.777187 |
| 6 | 7 | 'hbr' | 'group' | -0.3924697 | -0.3848339 | 0.70115395 | 0.87367661 |
| 6 | 7 | 'hbr' | 'b_AVD:grou | 1.67381344 | 1.27589856 | 0.20486117 | 0.52281149 |
| 6 | 7 | 'hbr' | 'c_V:group' | 2.17721831 | 1.65977952 | 0.10000055 | 0.37037242 |

| Source | Detector | Signal type | Contrast | Beta | T-stat | P-value | Q-value |
|---|---|---|---|---|---|---|---|
| 7 | 6 | 'hbo' | 'b_AVD' | 0.34600145 | 0.52298702 | 0.60210653 | 0.80717229 |
| 7 | 6 | 'hbo' | 'c_V' | 0.26854259 | 0.40189644 | 0.68859297 | 0.86888703 |
| 7 | 6 | 'hbo' | 'group' | -0.4439327 | -0.3367335 | 0.73700291 | 0.87738441 |
| 7 | 6 | 'hbo' | 'b_AVD:grou | -0.3726533 | -0.2243035 | 0.8229655 | 0.93518807 |
| 7 | 6 | 'hbo' | 'c_V:group' | -1.178441 | -0.7054037 | 0.48215133 | 0.72777559 |
| 7 | 6 | 'hbr' | 'b_AVD' | 0.74991086 | 1.59786711 | 0.11313596 | 0.39140779 |
| 7 | 6 | 'hbr' | 'c_V' | 0.04524995 | 0.09628209 | 0.92348362 | 0.96699856 |
| 7 | 6 | 'hbr' | 'group' | 0.61228002 | 0.65930245 | 0.51117294 | 0.74472351 |
| 7 | 6 | 'hbr' | 'b_AVD:grou | 1.56168119 | 1.32702357 | 0.18743515 | 0.50658148 |
| 7 | 6 | 'hbr' | 'c_V:group' | -1.7911729 | -1.5246872 | 0.13040183 | 0.41397406 |
| 7 | 7 | 'hbo' | 'b_AVD' | 0.87255093 | 0.94651104 | 0.34610325 | 0.64391303 |
| 7 | 7 | 'hbo' | 'c_V' | -0.0063683 | -0.0067766 | 0.99460622 | 0.99709897 |
| 7 | 7 | 'hbo' | 'group' | 3.52013784 | 2.05605874 | **0.04230563** | 0.25090528 |
| 7 | 7 | 'hbo' | 'b_AVD:grou | -4.7117456 | -2.0332478 | **0.0445994** | 0.25090528 |
| 7 | 7 | 'hbo' | 'c_V:group' | -3.9876667 | -1.6990057 | 0.09233671 | 0.35175888 |
| 7 | 7 | 'hbr' | 'b_AVD' | 1.44036695 | 2.28918021 | **0.0241094** | 0.19445821 |
| 7 | 7 | 'hbr' | 'c_V' | 2.57912576 | 4.04526184 | **0.00010117** | 0.01011726 |
| 7 | 7 | 'hbr' | 'group' | -0.5958408 | -0.504916 | 0.61469644 | 0.81148044 |
| 7 | 7 | 'hbr' | 'b_AVD:grou | 3.11595228 | 1.99370247 | **0.04882835** | 0.25699131 |
| 7 | 7 | 'hbr' | 'c_V:group' | 3.47984522 | 2.20570034 | **0.02962749** | 0.2116249 |
| 7 | 8 | 'hbo' | 'b_AVD' | 0.66921732 | 0.71276949 | 0.47760006 | 0.72777559 |
| 7 | 8 | 'hbo' | 'c_V' | 0.62244016 | 0.65003367 | 0.51711767 | 0.74472351 |
| 7 | 8 | 'hbo' | 'group' | 0.66902981 | 0.35901651 | 0.72031764 | 0.87399027 |
| 7 | 8 | 'hbo' | 'b_AVD:grou | -1.491908 | -0.5988896 | 0.55056141 | 0.76202271 |
| 7 | 8 | 'hbo' | 'c_V:group' | 0.29742888 | 0.11801018 | 0.90628942 | 0.9636272 |
| 7 | 8 | 'hbr' | 'b_AVD' | 0.82320354 | 1.41504339 | 0.16007204 | 0.46784048 |
| 7 | 8 | 'hbr' | 'c_V' | 0.45601762 | 0.77357108 | 0.4409563 | 0.71417152 |
| 7 | 8 | 'hbr' | 'group' | -0.5563633 | -0.4837744 | 0.6295729 | 0.82029042 |
| 7 | 8 | 'hbr' | 'b_AVD:grou | 1.95306765 | 1.28603935 | 0.20131285 | 0.52156971 |
| 7 | 8 | 'hbr' | 'c_V:group' | 1.88355308 | 1.23411104 | 0.21996996 | 0.54026705 |
| 8 | 7 | 'hbo' | 'b_AVD' | 0.10458816 | 0.14373289 | 0.88599223 | 0.95949752 |
| 8 | 7 | 'hbo' | 'c_V' | 1.03055681 | 1.38747495 | 0.16829162 | 0.4808332 |
| 8 | 7 | 'hbo' | 'group' | 2.51420233 | 1.74148485 | 0.08458423 | 0.34536956 |
| 8 | 7 | 'hbo' | 'b_AVD:grou | -2.9109099 | -1.5528066 | 0.1235367 | 0.40589047 |
| 8 | 7 | 'hbo' | 'c_V:group' | -1.348318 | -0.7073019 | 0.48097615 | 0.72777559 |
| 8 | 7 | 'hbr' | 'b_AVD' | -0.6250083 | -1.2856766 | 0.20143899 | 0.52156971 |
| 8 | 7 | 'hbr' | 'c_V' | 0.04052133 | 0.08202488 | 0.93478612 | 0.97581672 |
| 8 | 7 | 'hbr' | 'group' | -1.2893678 | -1.3295503 | 0.1866036 | 0.50658148 |
| 8 | 7 | 'hbr' | 'b_AVD:grou | 1.38467968 | 1.11069223 | 0.26928697 | 0.58842528 |
| 8 | 7 | 'hbr' | 'c_V:group' | 2.52588719 | 2.01656232 | **0.04634391** | 0.25090528 |
| 8 | 8 | 'hbo' | 'b_AVD' | -0.5758012 | -0.7110515 | 0.47865944 | 0.72777559 |
| 8 | 8 | 'hbo' | 'c_V' | 2.64008781 | 3.20367013 | **0.00180619** | 0.0436813 |
| 8 | 8 | 'hbo' | 'group' | -2.203647 | -1.3743238 | 0.17232419 | 0.48542025 |
| 8 | 8 | 'hbo' | 'b_AVD:grou | 2.57872974 | 1.25036862 | 0.21399812 | 0.54026705 |
| 8 | 8 | 'hbo' | 'c_V:group' | 1.7881622 | 0.85318063 | 0.39553886 | 0.7046095 |
| 8 | 8 | 'hbr' | 'b_AVD' | -0.5498277 | -1.1890094 | 0.23716994 | 0.55646806 |
| 8 | 8 | 'hbr' | 'c_V' | 0.15015667 | 0.31801808 | 0.75111509 | 0.8888936 |
| 8 | 8 | 'hbr' | 'group' | -0.7476871 | -0.8196687 | 0.4142977 | 0.71004527 |
| 8 | 8 | 'hbr' | 'b_AVD:grou | 3.17212199 | 2.69866757 | **0.00813741** | 0.10499888 |
| 8 | 8 | 'hbr' | 'c_V:group' | 2.04316466 | 1.721692 | 0.0881273 | 0.34536956 |

| Source | Detector | Signal type | Contrast | Beta | T-stat | P-value | Q-value |
|---|---|---|---|---|---|---|---|
| 9 | 9 | 'hbo' | 'b_AVD' | 0.03737245 | 0.05718498 | 0.95450863 | 0.9768626 |
| 9 | 9 | 'hbo' | 'c_V' | -0.388184 | -0.5927954 | 0.5546174 | 0.76498952 |
| 9 | 9 | 'hbo' | 'group' | 0.47365702 | 0.36956061 | 0.71246861 | 0.87399027 |
| 9 | 9 | 'hbo' | 'b_AVD:grou | -0.8557112 | -0.5150973 | 0.60758872 | 0.80742687 |
| 9 | 9 | 'hbo' | 'c_V:group' | -3.5722734 | -2.1615004 | **0.03297309** | 0.22740065 |
| 9 | 9 | 'hbr' | 'b_AVD' | -0.1473253 | -0.443228 | 0.6585304 | 0.84036742 |
| 9 | 9 | 'hbr' | 'c_V' | 0.76114412 | 2.28590942 | **0.02430728** | 0.19445821 |
| 9 | 9 | 'hbr' | 'group' | -0.3288175 | -0.477455 | 0.63404976 | 0.82077638 |
| 9 | 9 | 'hbr' | 'b_AVD:grou | -0.4455936 | -0.5608759 | 0.57610025 | 0.78380986 |
| 9 | 9 | 'hbr' | 'c_V:group' | 2.68657981 | 3.38065898 | **0.00102266** | 0.02921893 |
| 9 | 10 | 'hbo' | 'b_AVD' | -0.8777592 | -1.111995 | 0.26872906 | 0.58842528 |
| 9 | 10 | 'hbo' | 'c_V' | 0.51575366 | 0.64642082 | 0.51944465 | 0.74472351 |
| 9 | 10 | 'hbo' | 'group' | 0.24886156 | 0.17121444 | 0.86439106 | 0.95949752 |
| 9 | 10 | 'hbo' | 'b_AVD:grou | -1.0910738 | -0.5533808 | 0.58120193 | 0.78807042 |
| 9 | 10 | 'hbo' | 'c_V:group' | -0.8810037 | -0.4456447 | 0.65678923 | 0.84036742 |
| 9 | 10 | 'hbr' | 'b_AVD' | 0.83749845 | 1.94764899 | 0.0541792 | 0.27432505 |
| 9 | 10 | 'hbr' | 'c_V' | 0.03299736 | 0.0759963 | 0.93956946 | 0.97617606 |
| 9 | 10 | 'hbr' | 'group' | 0.12117223 | 0.14322079 | 0.88639563 | 0.95949752 |
| 9 | 10 | 'hbr' | 'b_AVD:grou | 1.10461205 | 1.03358669 | 0.30375015 | 0.60586223 |
| 9 | 10 | 'hbr' | 'c_V:group' | 1.10052675 | 1.02884947 | 0.30596042 | 0.60586223 |
| 10 | 9 | 'hbo' | 'b_AVD' | -0.9397548 | -1.4924528 | 0.13863722 | 0.4293322 |
| 10 | 9 | 'hbo' | 'c_V' | 0.70333543 | 1.10745774 | 0.27067563 | 0.58842528 |
| 10 | 9 | 'hbo' | 'group' | -0.746255 | -0.6091947 | 0.54373682 | 0.7604227 |
| 10 | 9 | 'hbo' | 'b_AVD:grou | 1.34041652 | 0.83542461 | 0.4054124 | 0.70655958 |
| 10 | 9 | 'hbo' | 'c_V:group' | 0.78559515 | 0.48698147 | 0.6273062 | 0.82000811 |
| 10 | 9 | 'hbr' | 'b_AVD' | 0.67863391 | 2.1958131 | **0.03034901** | 0.21297552 |
| 10 | 9 | 'hbr' | 'c_V' | -0.0183325 | -0.0587677 | 0.95325102 | 0.9768626 |
| 10 | 9 | 'hbr' | 'group' | -0.4612276 | -0.7340464 | 0.46458814 | 0.72777559 |
| 10 | 9 | 'hbr' | 'b_AVD:grou | 2.78936471 | 3.68643758 | **0.00036489** | 0.01582182 |
| 10 | 9 | 'hbr' | 'c_V:group' | 1.37511029 | 1.80700302 | 0.07368126 | 0.32747228 |
| 10 | 10 | 'hbo' | 'b_AVD' | -1.6210463 | -3.0487835 | **0.00292043** | 0.06142127 |
| 10 | 10 | 'hbo' | 'c_V' | -0.4265204 | -0.8006894 | 0.42515455 | 0.71155574 |
| 10 | 10 | 'hbo' | 'group' | -1.2725227 | -1.2749271 | 0.20520351 | 0.52281149 |
| 10 | 10 | 'hbo' | 'b_AVD:grou | -0.2430969 | -0.1837587 | 0.85456404 | 0.95790267 |
| 10 | 10 | 'hbo' | 'c_V:group' | 1.11748385 | 0.84884153 | 0.39793798 | 0.7046095 |
| 10 | 10 | 'hbr' | 'b_AVD' | 0.78838887 | 2.70613479 | **0.00796874** | 0.10499888 |
| 10 | 10 | 'hbr' | 'c_V' | 0.35112196 | 1.20212939 | 0.23206987 | 0.55254732 |
| 10 | 10 | 'hbr' | 'group' | -0.1433647 | -0.2380564 | 0.8123103 | 0.93266399 |
| 10 | 10 | 'hbr' | 'b_AVD:grou | 1.05170001 | 1.4798635 | 0.14196188 | 0.4334714 |
| 10 | 10 | 'hbr' | 'c_V:group' | -0.2728845 | -0.3830164 | 0.70249691 | 0.87367661 |
| 10 | 11 | 'hbo' | 'b_AVD' | 0.27021278 | 0.34307012 | 0.73224492 | 0.87613134 |
| 10 | 11 | 'hbo' | 'c_V' | 0.53247968 | 0.66776295 | 0.50577841 | 0.74472351 |
| 10 | 11 | 'hbo' | 'group' | -1.6632759 | -1.0890635 | 0.27866743 | 0.59258348 |
| 10 | 11 | 'hbo' | 'b_AVD:grou | -1.651624 | -0.8157758 | 0.41651092 | 0.71004527 |
| 10 | 11 | 'hbo' | 'c_V:group' | -2.8152356 | -1.3775042 | 0.17134232 | 0.48542025 |
| 10 | 11 | 'hbr' | 'b_AVD' | 0.01335066 | 0.0368682 | 0.97066145 | 0.98795059 |
| 10 | 11 | 'hbr' | 'c_V' | -0.3022351 | -0.8307469 | 0.40803816 | 0.70655958 |
| 10 | 11 | 'hbr' | 'group' | -0.0775965 | -0.1055949 | 0.91610912 | 0.96432539 |
| 10 | 11 | 'hbr' | 'b_AVD:grou | 0.7434219 | 0.8485412 | 0.39810437 | 0.7046095 |
| 10 | 11 | 'hbr' | 'c_V:group' | 1.25379582 | 1.4341012 | 0.1545724 | 0.4579923 |

| Source | Detector | Signal type | Contrast | Beta | T-stat | P-value | Q-value |
|---|---|---|---|---|---|---|---|
| 11 | 10 | 'hbo' | 'b_AVD' | -1.8210934 | -2.6464479 | **0.0094105** | 0.10997458 |
| 11 | 10 | 'hbo' | 'c_V' | -0.9908218 | -1.4344916 | 0.15446129 | 0.4579923 |
| 11 | 10 | 'hbo' | 'group' | -1.7176123 | -1.3150415 | 0.19141618 | 0.51386893 |
| 11 | 10 | 'hbo' | 'b_AVD:grou | -1.2786535 | -0.7309932 | 0.46644294 | 0.72777559 |
| 11 | 10 | 'hbo' | 'c_V:group' | -0.2518607 | -0.1439414 | 0.88582799 | 0.95949752 |
| 11 | 10 | 'hbr' | 'b_AVD' | 1.11086809 | 2.84712435 | **0.00532565** | 0.08199133 |
| 11 | 10 | 'hbr' | 'c_V' | -0.1836278 | -0.4687851 | 0.64021395 | 0.82608251 |
| 11 | 10 | 'hbr' | 'group' | 2.80495363 | 3.63884842 | **0.00043003** | 0.01582182 |
| 11 | 10 | 'hbr' | 'b_AVD:grou | -1.8307726 | -1.8716192 | 0.0640968 | 0.30276266 |
| 11 | 10 | 'hbr' | 'c_V:group' | -1.6838998 | -1.7172841 | 0.08893266 | 0.34536956 |
| 11 | 11 | 'hbo' | 'b_AVD' | 0.13042967 | 0.16879534 | 0.86628866 | 0.95949752 |
| 11 | 11 | 'hbo' | 'c_V' | -0.2180567 | -0.2811857 | 0.77913207 | 0.91662597 |
| 11 | 11 | 'hbo' | 'group' | -0.8523595 | -0.5665844 | 0.57222912 | 0.78120016 |
| 11 | 11 | 'hbo' | 'b_AVD:grou | -1.5435427 | -0.7735003 | 0.44099801 | 0.71417152 |
| 11 | 11 | 'hbo' | 'c_V:group' | -0.1940011 | -0.0968424 | 0.92303971 | 0.96699856 |
| 11 | 11 | 'hbr' | 'b_AVD' | -0.1347573 | -0.3510865 | 0.72624059 | 0.87613134 |
| 11 | 11 | 'hbr' | 'c_V' | 0.28483385 | 0.73897202 | 0.46160467 | 0.72777559 |
| 11 | 11 | 'hbr' | 'group' | 0.83463559 | 1.06576015 | 0.28902397 | 0.59258348 |
| 11 | 11 | 'hbr' | 'b_AVD:grou | -1.0120502 | -1.0726369 | 0.28594077 | 0.59258348 |
| 11 | 11 | 'hbr' | 'c_V:group' | -1.7853757 | -1.8789816 | 0.06307419 | 0.30276266 |
| 11 | 12 | 'hbo' | 'b_AVD' | -0.6591631 | -0.8363052 | 0.40491925 | 0.70655958 |
| 11 | 12 | 'hbo' | 'c_V' | 1.03799754 | 1.30298326 | 0.19548591 | 0.51963913 |
| 11 | 12 | 'hbo' | 'group' | -3.8797605 | -2.6383708 | **0.00962278** | 0.10997458 |
| 11 | 12 | 'hbo' | 'b_AVD:grou | -0.7202422 | -0.3666379 | 0.71464123 | 0.87399027 |
| 11 | 12 | 'hbo' | 'c_V:group' | 4.07259148 | 2.05107139 | **0.0427983** | 0.25090528 |
| 11 | 12 | 'hbr' | 'b_AVD' | 0.08151959 | 0.21706224 | 0.82858922 | 0.93891129 |
| 11 | 12 | 'hbr' | 'c_V' | -0.1614794 | -0.425537 | 0.67133309 | 0.85248646 |
| 11 | 12 | 'hbr' | 'group' | 0.15351753 | 0.20319493 | 0.83938355 | 0.94845599 |
| 11 | 12 | 'hbr' | 'b_AVD:grou | -0.3494042 | -0.3789651 | 0.70549386 | 0.87367661 |
| 11 | 12 | 'hbr' | 'c_V:group' | -0.6440303 | -0.6905817 | 0.49138174 | 0.73352882 |
| 12 | 11 | 'hbo' | 'b_AVD' | 0.99613358 | 1.52627504 | 0.13000633 | 0.41397406 |
| 12 | 11 | 'hbo' | 'c_V' | 0.04595219 | 0.07035945 | 0.94404399 | 0.97642763 |
| 12 | 11 | 'hbo' | 'group' | 0.27126562 | 0.22502143 | 0.82240844 | 0.93518807 |
| 12 | 11 | 'hbo' | 'b_AVD:grou | -3.2935525 | -2.0891377 | **0.0391596** | 0.24863239 |
| 12 | 11 | 'hbo' | 'c_V:group' | 0.86521061 | 0.5474952 | 0.58522299 | 0.79084187 |
| 12 | 11 | 'hbr' | 'b_AVD' | 0.47246281 | 1.36978312 | 0.1737334 | 0.48596756 |
| 12 | 11 | 'hbr' | 'c_V' | 0.6025984 | 1.73472569 | 0.08578079 | 0.34536956 |
| 12 | 11 | 'hbr' | 'group' | -0.096021 | -0.1361114 | 0.89199886 | 0.96172384 |
| 12 | 11 | 'hbr' | 'b_AVD:grou | 0.73978262 | 0.87611827 | 0.38300442 | 0.69009805 |
| 12 | 11 | 'hbr' | 'c_V:group' | 0.00751921 | 0.00882905 | 0.99297261 | 0.99709897 |
| 12 | 12 | 'hbo' | 'b_AVD' | -0.4283884 | -0.4777033 | 0.6338736 | 0.82077638 |
| 12 | 12 | 'hbo' | 'c_V' | 0.6980912 | 0.77717969 | 0.43883408 | 0.71417152 |
| 12 | 12 | 'hbo' | 'group' | -3.2499821 | -1.8699037 | 0.06433707 | 0.30276266 |
| 12 | 12 | 'hbo' | 'b_AVD:grou | 0.97566752 | 0.44162213 | 0.65968842 | 0.84036742 |
| 12 | 12 | 'hbo' | 'c_V:group' | 0.76067407 | 0.34223727 | 0.73286969 | 0.87613134 |
| 12 | 12 | 'hbr' | 'b_AVD' | 1.51667151 | 3.19496115 | **0.00185646** | 0.0436813 |
| 12 | 12 | 'hbr' | 'c_V' | 0.42556847 | 0.88514327 | 0.37814108 | 0.68441824 |
| 12 | 12 | 'hbr' | 'group' | -0.1366564 | -0.1417743 | 0.88753521 | 0.95949752 |
| 12 | 12 | 'hbr' | 'b_AVD:grou | 0.97506752 | 0.83317273 | 0.40667519 | 0.70655958 |
| 12 | 12 | 'hbr' | 'c_V:group' | -1.2214757 | -1.0329188 | 0.30406114 | 0.60586223 |

| Source | Detector | Signal type | Contrast | Beta | T-stat | P-value | Q-value |
|---|---|---|---|---|---|---|---|
| 12 | 13 | 'hbo' | 'b_AVD' | -0.757409 | -0.6975347 | 0.48703979 | 0.73239065 |
| 12 | 13 | 'hbo' | 'c_V' | 0.06143701 | 0.05671363 | 0.95488319 | 0.9768626 |
| 12 | 13 | 'hbo' | 'group' | 0.32076255 | 0.15345159 | 0.8783424 | 0.95949752 |
| 12 | 13 | 'hbo' | 'b_AVD:grou | -3.3139434 | -1.1543089 | 0.25104477 | 0.5705563 |
| 12 | 13 | 'hbo' | 'c_V:group' | -0.3141474 | -0.1094775 | 0.91303677 | 0.9636272 |
| 12 | 13 | 'hbr' | 'b_AVD' | 1.01573431 | 1.92301109 | 0.05724029 | 0.28266808 |
| 12 | 13 | 'hbr' | 'c_V' | 0.69061592 | 1.28514954 | 0.20162237 | 0.52156971 |
| 12 | 13 | 'hbr' | 'group' | 1.24724532 | 1.1868025 | 0.23803564 | 0.55646806 |
| 12 | 13 | 'hbr' | 'b_AVD:grou | 1.42129302 | 1.06541572 | 0.28917899 | 0.59258348 |
| 12 | 13 | 'hbr' | 'c_V:group' | -0.5343727 | -0.3951481 | 0.69355072 | 0.87239084 |
| 13 | 12 | 'hbo' | 'b_AVD' | -0.785153 | -0.9575605 | 0.34052667 | 0.64250315 |
| 13 | 12 | 'hbo' | 'c_V' | 1.30894092 | 1.57850346 | 0.11751634 | 0.39501289 |
| 13 | 12 | 'hbo' | 'group' | -3.5724272 | -2.2352264 | **0.02756117** | 0.20800881 |
| 13 | 12 | 'hbo' | 'b_AVD:grou | -0.3732719 | -0.1771766 | 0.85971764 | 0.95790267 |
| 13 | 12 | 'hbo' | 'c_V:group' | -3.5113464 | -1.654374 | 0.10109592 | 0.3709942 |
| 13 | 12 | 'hbr' | 'b_AVD' | 0.38434786 | 1.03572524 | 0.30275589 | 0.60586223 |
| 13 | 12 | 'hbr' | 'c_V' | -0.3084049 | -0.821731 | 0.41312808 | 0.71004527 |
| 13 | 12 | 'hbr' | 'group' | 0.20674006 | 0.2749411 | 0.78391224 | 0.91954515 |
| 13 | 12 | 'hbr' | 'b_AVD:grou | -0.717087 | -0.7798509 | 0.437267 | 0.71417152 |
| 13 | 12 | 'hbr' | 'c_V:group' | -2.6592452 | -2.8706976 | **0.00497176** | 0.08199133 |
| 13 | 13 | 'hbo' | 'b_AVD' | -2.2055988 | -2.407104 | **0.01785959** | 0.17502023 |
| 13 | 13 | 'hbo' | 'c_V' | 1.70254228 | 1.85203813 | 0.06688433 | 0.30933131 |
| 13 | 13 | 'hbo' | 'group' | -2.1960581 | -1.2138344 | 0.22758692 | 0.54840223 |
| 13 | 13 | 'hbo' | 'b_AVD:grou | -0.3551099 | -0.1490317 | 0.88182005 | 0.95949752 |
| 13 | 13 | 'hbo' | 'c_V:group' | 0.77933215 | 0.32691985 | 0.74439194 | 0.88355126 |
| 13 | 13 | 'hbr' | 'b_AVD' | 0.33510716 | 0.75022079 | 0.45483207 | 0.72483198 |
| 13 | 13 | 'hbr' | 'c_V' | 0.1694546 | 0.37537386 | 0.70815434 | 0.87399027 |
| 13 | 13 | 'hbr' | 'group' | -1.8134541 | -2.0243677 | **0.04552072** | 0.25090528 |
| 13 | 13 | 'hbr' | 'b_AVD:grou | 2.17001323 | 1.92999097 | 0.05635861 | 0.28179306 |
| 13 | 13 | 'hbr' | 'c_V:group' | 0.70774498 | 0.62440987 | 0.5337391 | 0.75174522 |
| 14 | 14 | 'hbo' | 'b_AVD' | 0.21202991 | 0.12451123 | 0.90115319 | 0.9636272 |
| 14 | 14 | 'hbo' | 'c_V' | -3.8756644 | -2.2873903 | **0.02421751** | 0.19445821 |
| 14 | 14 | 'hbo' | 'group' | -0.5774242 | -0.1772678 | 0.85964615 | 0.95790267 |
| 14 | 14 | 'hbo' | 'b_AVD:grou | 0.58393232 | 0.12899358 | 0.89761428 | 0.96258904 |
| 14 | 14 | 'hbo' | 'c_V:group' | -7.1324125 | -1.5827033 | 0.11655496 | 0.39501289 |
| 14 | 14 | 'hbr' | 'b_AVD' | 1.62109437 | 2.09440856 | **0.0386774** | 0.24863239 |
| 14 | 14 | 'hbr' | 'c_V' | 0.60173165 | 0.77552414 | 0.43980697 | 0.71417152 |
| 14 | 14 | 'hbr' | 'group' | -1.9961088 | -1.3282414 | 0.18703403 | 0.50658148 |
| 14 | 14 | 'hbr' | 'b_AVD:grou | 3.28577819 | 1.63277802 | 0.10556916 | 0.38388784 |
| 14 | 14 | 'hbr' | 'c_V:group' | 4.08845622 | 2.02202189 | **0.04576679** | 0.25090528 |
| 14 | 15 | 'hbo' | 'b_AVD' | -1.0089181 | -0.9384387 | 0.35021444 | 0.64647014 |
| 14 | 15 | 'hbo' | 'c_V' | -1.0699027 | -0.9779727 | 0.33037927 | 0.63596069 |
| 14 | 15 | 'hbo' | 'group' | 6.51512323 | 3.03225985 | **0.00307106** | 0.06142127 |
| 14 | 15 | 'hbo' | 'b_AVD:grou | -6.3189847 | -2.225293 | **0.02824181** | 0.20919859 |
| 14 | 15 | 'hbo' | 'c_V:group' | -6.1570659 | -2.1451145 | **0.03429427** | 0.2286285 |
| 14 | 15 | 'hbr' | 'b_AVD' | -0.6193888 | -0.739192 | 0.46147167 | 0.72777559 |
| 14 | 15 | 'hbr' | 'c_V' | -0.0519694 | -0.0618064 | 0.9508368 | 0.9768626 |
| 14 | 15 | 'hbr' | 'group' | 1.24428036 | 0.79385098 | 0.42910731 | 0.71417152 |
| 14 | 15 | 'hbr' | 'b_AVD:grou | 0.31863473 | 0.14632154 | 0.88395356 | 0.95949752 |
| 14 | 15 | 'hbr' | 'c_V:group' | -1.3550739 | -0.6206198 | 0.53622065 | 0.75259038 |

| Source | Detector | Signal type | Contrast | Beta | T-stat | P-value | Q-value |
|---|---|---|---|---|---|---|---|
| 15 | 14 | 'hbo' | 'b_AVD' | -1.1680331 | -0.7054641 | 0.48211392 | 0.72777559 |
| 15 | 14 | 'hbo' | 'c_V' | -0.3015419 | -0.1786055 | 0.8585983 | 0.95790267 |
| 15 | 14 | 'hbo' | 'group' | 0.22546922 | 0.06954112 | 0.94469373 | 0.97642763 |
| 15 | 14 | 'hbo' | 'b_AVD:grou | -1.667399 | -0.3823415 | 0.70299584 | 0.87367661 |
| 15 | 14 | 'hbo' | 'c_V:group' | -0.0898566 | -0.0204041 | 0.9837605 | 0.99244316 |
| 15 | 14 | 'hbr' | 'b_AVD' | -0.4806141 | -0.5404051 | 0.59008435 | 0.7947264 |
| 15 | 14 | 'hbr' | 'c_V' | 0.72537506 | 0.80729886 | 0.42135468 | 0.71004527 |
| 15 | 14 | 'hbr' | 'group' | -2.8636697 | -1.808817 | 0.07339682 | 0.32747228 |
| 15 | 14 | 'hbr' | 'b_AVD:grou | 5.22979575 | 2.38941004 | **0.01869444** | 0.17502023 |
| 15 | 14 | 'hbr' | 'c_V:group' | 8.03390158 | 3.63544051 | **0.0004351** | 0.01582182 |
| 15 | 15 | 'hbo' | 'b_AVD' | 0.05142144 | 0.02943948 | 0.97657105 | 0.98893271 |
| 15 | 15 | 'hbo' | 'c_V' | -1.9094703 | -1.0808672 | 0.28228047 | 0.59258348 |
| 15 | 15 | 'hbo' | 'group' | -2.0873415 | -0.6245295 | 0.53366087 | 0.75174522 |
| 15 | 15 | 'hbo' | 'b_AVD:grou | 6.94121726 | 1.53906189 | 0.12685574 | 0.40921207 |
| 15 | 15 | 'hbo' | 'c_V:group' | -3.6932096 | -0.813381 | 0.41787589 | 0.71004527 |
| 15 | 15 | 'hbr' | 'b_AVD' | 0.61970791 | 0.64851706 | 0.51809382 | 0.74472351 |
| 15 | 15 | 'hbr' | 'c_V' | -0.7470451 | -0.7729913 | 0.44129781 | 0.71417152 |
| 15 | 15 | 'hbr' | 'group' | -0.5006246 | -0.2676027 | 0.78954028 | 0.92343892 |
| 15 | 15 | 'hbr' | 'b_AVD:grou | 4.30838026 | 1.723628 | 0.08777547 | 0.34536956 |
| 15 | 15 | 'hbr' | 'c_V:group' | 0.86037946 | 0.34105088 | 0.73375999 | 0.87613134 |
| 15 | 16 | 'hbo' | 'b_AVD' | 1.60029391 | 0.91748703 | 0.36103085 | 0.66244193 |
| 15 | 16 | 'hbo' | 'c_V' | -0.8695212 | -0.4941078 | 0.62228219 | 0.8161078 |
| 15 | 16 | 'hbo' | 'group' | 3.2990623 | 0.98845998 | 0.32524379 | 0.63596069 |
| 15 | 16 | 'hbo' | 'b_AVD:grou | -5.2164059 | -1.1364113 | 0.25842156 | 0.58400354 |
| 15 | 16 | 'hbo' | 'c_V:group' | -4.5294001 | -0.9785178 | 0.33011103 | 0.63596069 |
| 15 | 16 | 'hbr' | 'b_AVD' | 1.3605996 | 1.41448517 | 0.16023536 | 0.46784048 |
| 15 | 16 | 'hbr' | 'c_V' | 1.2151416 | 1.24543428 | 0.21579796 | 0.54026705 |
| 15 | 16 | 'hbr' | 'group' | 2.25343214 | 1.18769874 | 0.23768379 | 0.55646806 |
| 15 | 16 | 'hbr' | 'b_AVD:grou | -2.3150252 | -0.8999525 | 0.37024484 | 0.67624627 |
| 15 | 16 | 'hbr' | 'c_V:group' | 1.71001803 | 0.65884765 | 0.51146379 | 0.74472351 |
| 16 | 15 | 'hbo' | 'b_AVD' | 3.41819161 | 2.65280701 | **0.00924637** | 0.10997458 |
| 16 | 15 | 'hbo' | 'c_V' | 0.14226956 | 0.1094834 | 0.91303211 | 0.9636272 |
| 16 | 15 | 'hbo' | 'group' | 2.38553387 | 0.96010261 | 0.33925198 | 0.64250315 |
| 16 | 15 | 'hbo' | 'b_AVD:grou | -1.7307093 | -0.517951 | 0.60560321 | 0.80742687 |
| 16 | 15 | 'hbo' | 'c_V:group' | 0.53932691 | 0.16042938 | 0.87285707 | 0.95949752 |
| 16 | 15 | 'hbr' | 'b_AVD' | 0.17721436 | 0.26438626 | 0.7920106 | 0.92362753 |
| 16 | 15 | 'hbr' | 'c_V' | 0.71823259 | 1.07224549 | 0.28611566 | 0.59258348 |
| 16 | 15 | 'hbr' | 'group' | 0.93636367 | 0.8672187 | 0.38783806 | 0.69567366 |
| 16 | 15 | 'hbr' | 'b_AVD:grou | -0.2600174 | -0.1808115 | 0.85687084 | 0.95790267 |
| 16 | 15 | 'hbr' | 'c_V:group' | 2.10563897 | 1.48903795 | 0.13953296 | 0.4293322 |
| 16 | 16 | 'hbo' | 'b_AVD' | 2.84636843 | 2.07904766 | **0.04009715** | 0.2506072 |
| 16 | 16 | 'hbo' | 'c_V' | -0.7933195 | -0.5742798 | 0.56703058 | 0.777187 |
| 16 | 16 | 'hbo' | 'group' | -2.9760025 | -1.1699442 | 0.24472346 | 0.5593679 |
| 16 | 16 | 'hbo' | 'b_AVD:grou | 2.44093388 | 0.69044982 | 0.49146431 | 0.73352882 |
| 16 | 16 | 'hbo' | 'c_V:group' | -0.3906253 | -0.1098891 | 0.91271117 | 0.9636272 |
| 16 | 16 | 'hbr' | 'b_AVD' | 0.3364312 | 0.51063321 | 0.6107006 | 0.80887496 |
| 16 | 16 | 'hbr' | 'c_V' | 0.39929873 | 0.59930672 | 0.55028434 | 0.76202271 |
| 16 | 16 | 'hbr' | 'group' | 0.868936 | 0.66639346 | 0.50664954 | 0.74472351 |
| 16 | 16 | 'hbr' | 'b_AVD:grou | 0.21913721 | 0.13215058 | 0.895123 | 0.96249785 |
| 16 | 16 | 'hbr' | 'c_V:group' | -1.1392663 | -0.6802646 | 0.49786318 | 0.740317 |

#### Supplementary Material 7

Hemodynamic time-series were quantified using a deconvolution approach implemented in *nirs-toolbox* (Santosa et al., 2018). For each participant, HbO and HbR responses were modelled using a finite impulse response (FIR) approach with 1-s time bins spanning the 18-s stimulus period. Condition-specific time courses were obtained for each channel and subsequently averaged within ROIs and hemispheres. The time series are plotted by grouping optodes into regions of interest (ROIs) to provide a clearer representation of brain activity without implying that statistical inference was performed at the ROI level.

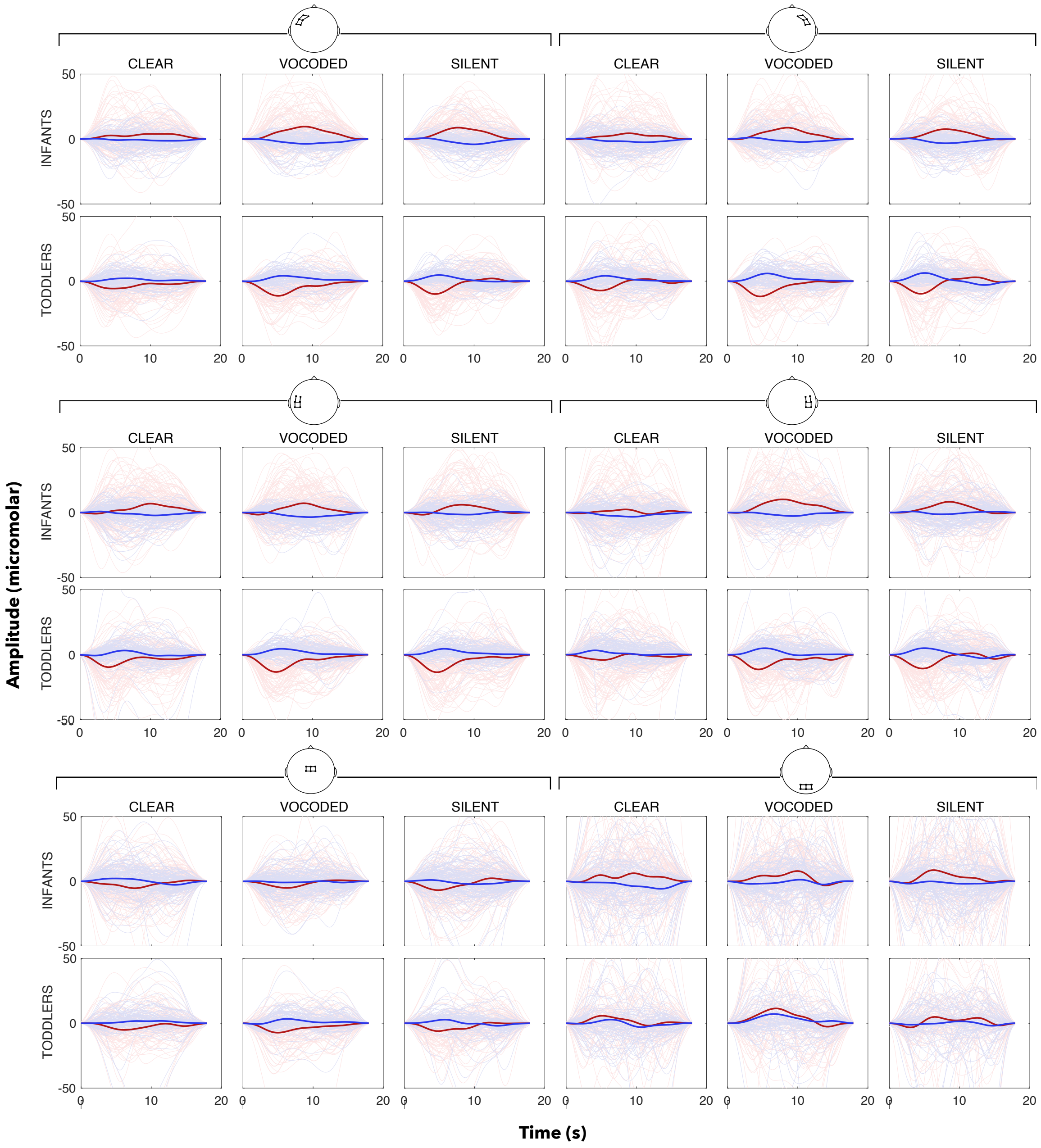

*Figure 5. Group-level hemodynamic responses (HbO, red; HbR, blue) for the infants and toddlers per condition averaged per ROI. The thick lines correspond to the mean value for all channels in each ROI across all participants in each group; the thin lines, the individual time series of each participant per channel in the ROI.*
